# Metabolic zonation and characterization of tissue slices with spatial transcriptomics

**DOI:** 10.1101/2025.11.11.687271

**Authors:** Oier Etxezarreta Arrastoa, Anna-Chiara Pirona, Allon Wagner, Nir Yosef

## Abstract

The exchanges of small molecules between cells and their environments are essential for the formation of functioning tissues. To study them at scale, we developed Harreman (Basque for “receive and give”), an algorithm for identifying metabolic crosstalk from spatially resolved transcriptomics of intact tissues. Unlike previous methods, which primarily focus on the secretion or reception of protein signals, Harreman reconstructs molecular metabolic crosstalk based on the co-localized expression of metabolite transporters. By utilizing a series of increasingly detailed models for testing spatial correlation, Harreman provides insight at multiple levels: a) coarse partition of the tissue into regions sharing metabolic characteristics; b) identification of metabolic exchanges within each region; and c) inference of the cell subsets involved in those exchanges. Harreman identified a sodium/calcium exchange at the tumor boundary in human lung metastases of human renal cancers, and associated it with nearby pro-inflammatory macrophages. In the mouse model of DSS-induced colitis, Harreman identified vitamin A and lysophosphatidylcholine transport at the interface of the epithelial monolayer as major signals associated with regeneration. Harreman is available at https://github.com/YosefLab/Harreman.

## Introduction

Spatially resolved transcriptomics has opened new avenues to study how cells interact within their native tissue context [1–3]. By providing gene expression profiles along with each cell’s spatial location, these technologies enable the identification of dependencies and communications between neighboring cells that were previously unattainable with dissociated single-cell data [4, 5]. Current analyses of cell–cell communication in such data have primarily focused on biochemical signaling via ligand–receptor (LR) interactions [6–13]. Indeed, a significant form of intercellular interaction is mediated by secreted ligands binding to receptors on adjacent cells, triggering downstream effects [14]. LR interaction databases containing LR pairs with their corresponding stimulators and inhibitors have been built [12]. However, this focus on protein-mediated signaling overlooks important roles of intercellular crosstalk via non-protein factors [15, 16]. Among these, metabolites and other small molecules function as important messengers between cells [17], a dimension of cellular communication that has remained comparatively underexplored in spatial transcriptomic analyses.

The exchange of nutrients and metabolites between cells and their surroundings can profoundly influence cellular behavior [18] and fate [19]. For example, in the tumor microenvironment, cancer cells often compete with immune cells for key nutrients to fuel their effector function [20]. Hypoxic regions in the tumor stimulate HIF-1*α* activity, thereby upregulating glycolysis and increasing the concentration of by-products in the environment, including lactate, which supports dysfunctional states in CD8+ T cells [21] and augments T regulatory phenotypes [22]. In the intestine, microbiota-derived metabolites are signals that mediate the crucial balance between immune tolerance to commensal microbes and immune rejection of pathogens. Microbiota-derived short-chain fatty acids (SCFAs), such as butyrate, support the differentiation of regulatory T cells (Tregs) [23, 24] and the induction of macrophage tolerance [25]. On the other hand, microbiota-derived SCFAs regulate B cell metabolism, supporting antibody production, thereby enhancing antibody response against pathogens [26]. Bile acid metabolites have also been reported to influence the differentiation of Tregs and Th17 cells [27].

Despite the importance of metabolite-driven cell interactions, most computational frameworks for cell–cell communication have yet to address this facet. Most methods are based on databases of ligand–receptor pairs that do not incorporate metabolite transporters and the molecules they carry, but some studies have already begun to address metabolite-mediated communication using transcriptomic data [28–33]. MEBOCOST [29] infers cell-cell communication based on metabolite-receptor interactions in single-cell RNA-seq data without considering the spatial position of cells. scCellFie [30] infers metabolic activities in single-cell and spatial transcriptomics and quantifies intercellular interactions by integrating metabolite-based ligand-receptor interaction information from MetalinksDB [28] with the metabolic activity and receptor gene expression in sender and receiver cells, respectively. This method uses a binary approach to compute the co-localization score instead of considering the continuous gene expression values, and it does not take the distance between cells in the neighborhood into account. Additionally, SpatialDM [34] is a spatially-weighted method that detects co-localized ligand-receptor pairs but does not consider metabolite information and cell type identity. Therefore, dedicated strategies to exploit spatial transcriptomics for inferring metabolic exchanges between neighboring cells considering the cell type identity as well as the deconvolved expression data from spot-based technologies are largely lacking. This gap highlights the need for new computational tools that can identify metabolite-based cell interactions directly from spatial omics data.

Here, we introduce Harreman (Basque for “receive and give”), a computational tool for studying metabolic crosstalk events in spatial transcriptomics. Harreman leverages spatial autocorrelation in gene expression at different levels: from tissue zonation into metabolically similar regions, to metabolite exchange among cells within each zone, and ultimately to identify the cell populations involved in those signaling events. Harreman introduces a set of statistical models to evaluate spatially-weighted correlation between the expression patterns of genes coding for metabolite transporters across the tissue, under the premise that if a metabolite is actively shared or present in a microenvironment, the cells exporting and importing that metabolite may show coordinated spatial expression of the corresponding genes. The algorithm supports cell-type-agnostic analysis, which does not require classification of cell types, as well as cell-type-aware analysis, which focuses on interactions of specific cell subsets. Harreman can also incorporate deconvolved spatial data from spot-based spatial transcriptomic assays. Harreman builds and generalizes our previously published Hotspot algorithm [35]. To accelerate Harreman’s runtime, we reimplemented the Hotspot codebase using PyTorch.

In addition to developing the Harreman algorithm, we introduce HarremanDB, a database of metabolite-transporter associations in human and mouse. We collected information on metabolites and their corresponding transporters from publicly available platforms [28, 29, 36–42], and added manually curated pairs from relevant research literature. Protein ligand-receptor interactions from CellChatDB [43] have also been incorporated.

We demonstrate Harreman’s approach in two relevant biological contexts: the tumor microenvironment and gut inflammation. In human lung metastases of renal cell carcinomas (RCCs), we found that a high abundance of sodium/calcium exchange through the *SLC8A1* transporter in the tumor boundary shapes macrophage function. In the inflamed colon, Harreman identified region- and condition-specific metabolic niches and metabolite signaling that may drive or temper inflammation. Further, metabolites such as vitamin A and lysophosphatidylcholine (a substrate of the *Mfsd2a* transporter) were differentially abundant in the distal region of the regenerating colon, revealing metabolic processes that may support tissue regeneration following barrier damage.

## Results

### Harreman infers metabolic zonation and crosstalk in spatial transcriptomics datasets

Harreman relies on several forms of spatial correlation to offer increasingly detailed interpretations of metabolic activity in tissue slices profiled with spatial transcriptomics (Fig. 1 and Supplementary Table 1). At the coarsest level, Harreman divides the tissue into regions, each with a characteristic set of metabolic functions. This partition depends on groups (modules) of metabolic enzymes that tend to be expressed by the same cell or by nearby cells (i.e., spatially correlated). This first level of analysis relies on the Hotspot algorithm [35], which we previously introduced, while re-implementing it to allow it to scale to data magnitudes of spatial transcriptomics (Fig. S1 and Methods; **Test statistics 1 and 2**). Harreman supports both single-cell and spot-based spatial transcriptomics (Fig. 1A). Formally, we define for every cell (or spot) a list of neighbors (using KNN or taking all neighbors up to a certain distance), weighted by their distance (Fig. 1B), and compute a spatial correlation statistic (all formulas are summarized in Supplementary Table 1 and Methods; Fig. 1C). For significance testing, we utilize a theoretical null model (following [35]), which we demonstrate to be consistent with an empirical test (shuffling the locations of cells; Fig. 1D, Fig. S1, and Fig. S2). At the next level, Harreman generates hypotheses about which metabolites are exchanged in the tissue or within each zone (Fig. 1C and Methods; **Test statistic 3**). This is done by calculating an aggregate spatial correlation, considering all pairs of genes that are involved in the import or export of the respective metabolite. Spatially co-localized metabolites can also be grouped together (Fig. 1C and Methods; **Test statistic 4**). Further increasing the level of detail, Harreman can also predict which subsets of cells are involved in the exchange of specific metabolic signals (Fig. 1C and Methods; **Test statistic 6**). This is done by an aggregate spatial correlation, considering cell-type restricted expression of pairs of import or export molecules defined in **Test statistic 5**. In addition to these core capacities, on which we focus in this paper, Harreman implements a series of aggregate spatial correlation statistics to identify different scenarios of interaction, e.g., testing whether a given metabolite is exchanged by cells of a certain type with all other cells in their neighborhood (Fig. 1C, Methods, and Supplementary Table 1; **Additional tests for interaction**).

**Figure 1.**
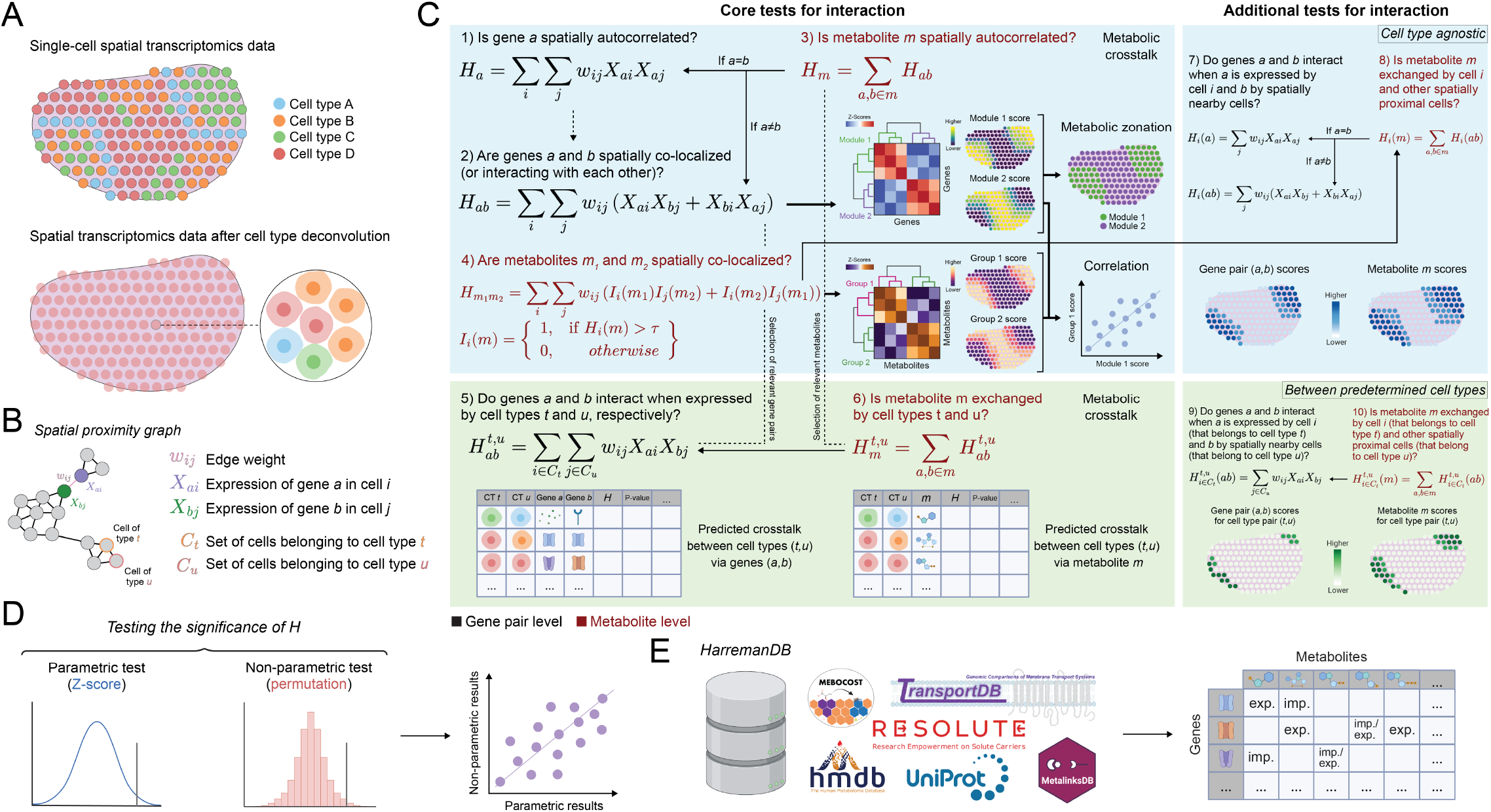
Overview of the Harreman algorithm. (A-B) The Harreman algorithm takes as input single-cell or spot-based spatial transcriptomics data (A) and creates a spatial neighborhood graph based on either nearest neighbors or spatial distance (B). (C) Metabolic zonation of the tissue is inferred from gene co-expression (spatially autocorrelated genes selected from test statistic 1 are used in test statistic 2) weighted by spatial proximity (1-2), and metabolic crosstalk inference within zones can be performed at the gene pair (colored in black) and the metabolite (colored in dark red) level by computing an aggregate spatial correlation, considering all pairs of genes that are involved in the import or export of the respective metabolite (1-3). Spatially co-localized metabolites can also be grouped together (4). Relevant gene pairs and/or metabolites can be selected to predict which cell types are involved in each communication event, either at the gene-pair or metabolite level (5-6). Apart from these core capacities, additional tests have been implemented to identify different interaction scenarios (7-10). The test statistics with a blue background do not require cell type information, while those with a green background use predetermined cell type information. Solid arrow from Y to X denotes that formula Y uses formula X; dashed arrows denote a possible influence; and solid thick arrows denote downstream analysis. CT: cell type; *m*: metabolite. (D) To assess statistical significance, a parametric approach and a non-parametric approach have been implemented, calculating the z-scores and building an empirical null distribution through permutations, respectively. (E) To infer metabolic crosstalk, we curated a metabolite transporter database for humans and mice by collecting metabolite and their corresponding transporter information from different publicly available platforms. imp.: importer; exp.: exporter. Some of the icons shown in this figure have been obtained from BioRender.

Harreman requires a priori knowledge of metabolic exchange reactions between cells and their environment. We curated a database of metabolites and their respective transporter genes from HMDB [36], MEBOCOST [29], TransportBD [37], MetalinksDB [28], RESOLUTE [38], augmented with manually curated metabolite-transporter information from the scientific literature (e.g., [39–42]). This effort resulted in HarremanDB - a database of metabolites and their associated transporters that consists of 1,047 human and 1,038 mouse metabolites. Two versions of the database were constructed: one including all metabolites regardless of subcellular localization, and another restricted to metabolites annotated as extracellular, based on information from resources such as HMDB (242 human and 249 mouse metabolites). All analyses in this study were performed using the extracellular version of the database (Fig. 1E and Methods).

### Metabolic zonation of lung metastases

We first applied Harreman to a dataset of lung metastasis originating from renal cell carcinoma (RCC) [44], consisting of seven tissue sections of the same patient, profiled with Slide-seq [45] (200,394 spots of 10 *µ*m in diameter in total; Fig. 2A and Methods). The first level of analysis provided by Harreman identified four modules of spatially colocalized enzyme-coding genes (Fig. 2B, Fig. S3, and Methods; **Test statistics 1 and 2**). These modules are present in all seven slides and are robust to changes in the neighborhood radius considered around each spot as well as the minimum number of genes selected for module computation (Fig. S4, Methods). The cumulative expression of genes within each module induces a partition of each slide into four respective metabolic zones (Fig. 2C). We interpreted the zones based on the associated histology images from [44]. The zone corresponding to Module 1 occupies most of the tumor area, while Module 2 sits at the boundary, and Modules 3 and 4 reside at peri-tumoral or adjacent-normal tissue areas.

**Figure 2.**
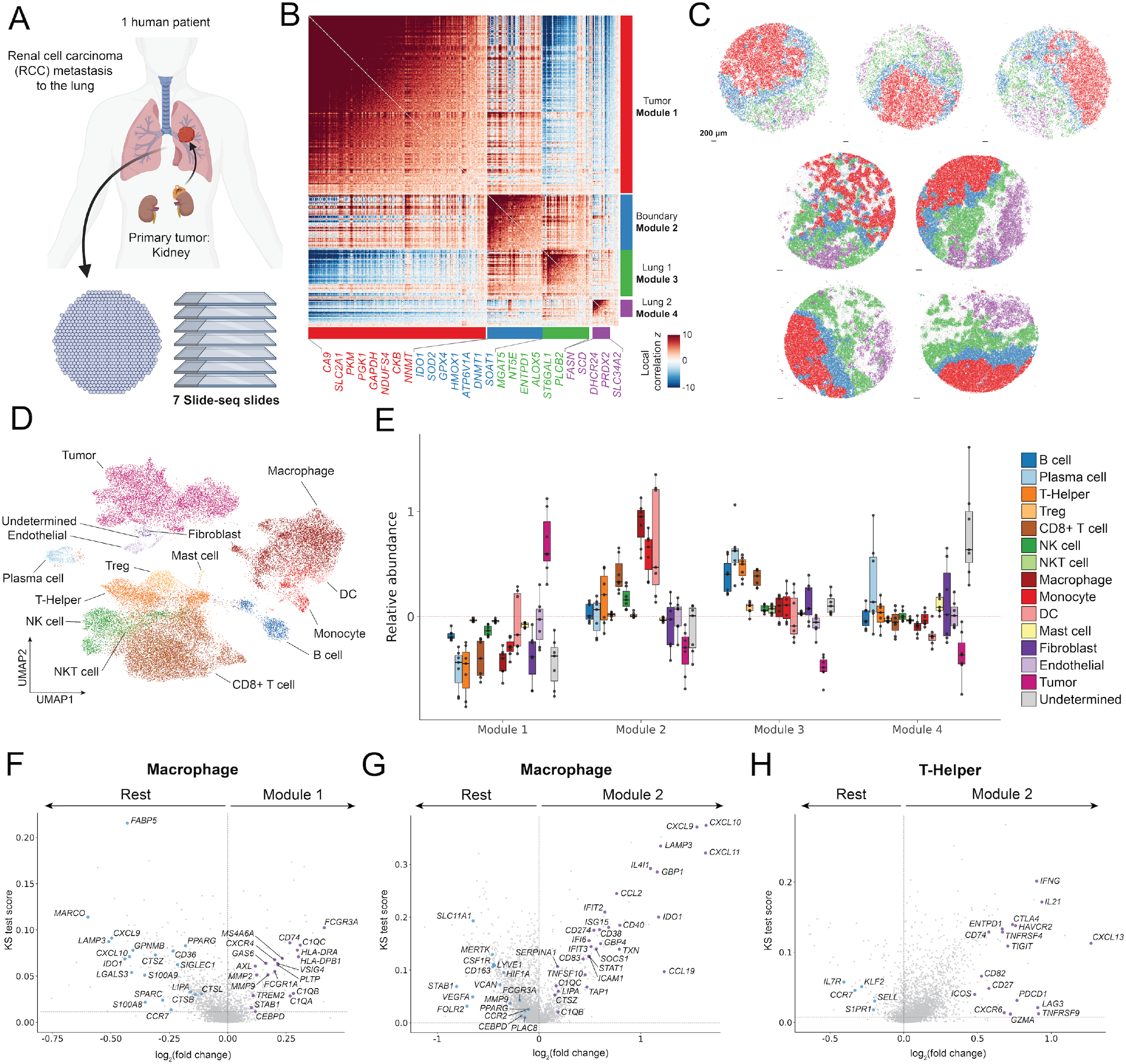
Metabolic characterization of lung metastases in renal cell carcinoma. (A) Schematic representation of the dataset [44], where 7 tissue sections from lung metastasis of a single human patient were profiled with Slide-seq [45]. Figure created using [BioRender](https://www.biorender.com/). (B) Cluster map of 605 metabolic genes that are significantly spatially autocorrelated (theoretical FDR < 0.01; **Test statistic 1**) shows four main modules (using **Test statistic 2**). (C) Spatial distribution of modules across samples. Scale bars, 200 *µ*m. (D) UMAP plot of the single-cell dataset [46] used as a reference for the cell type deconvolution analysis. Cell types are colored according to (E). (E) Box plot showing the relative abundance of each cell type across modules. Each dot represents a tissue slide. (F-H) Volcano plots showing the cell type-specific differential expression (DE) analysis between Module 1 and the rest in macrophages (F), between Module 2 and the rest in macrophages (G), and between Module 2 and the rest in T-Helper cells (H). Spots with a cell type proportion higher than 1 standard deviation above the mean for the corresponding cell type were selected for the DE analysis. Relevant genes are labeled. KS: Kolmogorov-Smirnov.

To explore how this metabolism-centric zonation reflects cellular composition and function, we analyzed the relative abundance of cell types across modules using deconvolution [47] with an annotated single-cell RNA-seq reference [46] (Fig. 2D) that was also used in the original study (Fig. 2E and Methods). In addition to supporting the enrichment of tumor cells in the malignant region delineated by Module 1, this analysis pointed to over-representation of myeloid cells (macrophages, monocytes, and dendritic cells) and the adaptive arm of the immune system in Modules 2 and 3, respectively (Fig. 2E, Methods). This pattern is consistent with current knowledge on the organization of immunity in solid tumors, with concentration of myeloid cells at the tumor boundary, supporting tumor invasion through matrix remodeling and angiogenesis [48, 49]. It is also consistent with the presence of normal residency of T cells and immune exclusion of T and plasma cells in the adjacent, peri-tumoral tissue [50, 51].

The tumor stroma (captured by Modules 1 and 2) plays a pivotal role in promoting tumor progression and treatment resistance [52–55]. Accumulating evidence suggests that metastatic cells may precondition their target distant sites by stromal remodeling and immune modulation even before major colonization [56, 57]. To study how these processes may be reflected in the Harreman-based zonation, we focused on macrophages. We performed differential expression analysis comparing macrophages from the tumor (Module 1) or from the tumor boundary (Module 2) to those captured in the remaining tissue region (Fig. 2F-G). Macrophages in the Module 1 area exhibited an immunosuppressive, tumor-promoting phenotype, with elevated expression of *MMP2* and *MMP9* (metalloproteinases involved in extracellular matrix degradation and tumor invasion) as well as *CXCR4*, which mediates cell migration and metastasis, and *AXL*, a receptor tyrosine kinase linked to poor prognosis [58–61] (Fig. 2F). Macrophages from the Module 2 area displayed a more immune-activating phenotype. These cells upregulated interferon response genes such as *STAT1, IFIT2*, and *TAP1* [62, 63], along with the chemokines *CXCL9, CXCL10*, and *CXCL11*, suggesting active immunity and recruitment of cytotoxic T-cells via the *CXCR3* axis [64, 65] (Fig. 2G). To further explore this link at the tumor boundary, we investigated the gene expression profile of T-helper cells in Module 2 compared to T-helpers from all other regions (Fig. 2H). Module 2 T-helpers had high expression of antitumor effector genes, such as *IFNG, IL21*, and *TNFRSF4 (OX40)* [66–68] and of immune checkpoint genes including *CTLA4, TIGIT*, and *LAG3*, associated with T cell exhaustion [69–71], consistent with antigen exposure and activation in the tumor site [72] (Fig. 2H). Indeed, T-helpers from the other modules had higher expression of genes associated with stemness, including *IL7R, SELL, CCR7*, and *KLF2* [73].

Next, we sought to characterize the metabolic activity within each region. In the following section, we turn back to Harreman and apply its additional modeling capacities (Supplementary Table 1) for a more detailed analysis of metabolic exchanges in each region, with a focus on the tumor boundary.

### Metabolite modules confirm conservation of RCC phenotype in lung metastasis

To better understand the metabolic activity across tumor regions, we applied Harreman in its metabolite discovery mode. Here, we aggregate the co-localization scores of all pairs of genes that are involved in the import or export of a given metabolite. We use those aggregate values to estimate the extent of the metabolite’s exchange and to identify the tissue regions in which it takes place (Supplementary Table 1 and Methods; **Test statistics 3 and 8**, respectively). In cases where a gene is designated as a transporter capable of both import and export, we include its spatial autocorrelation in our aggregation as well (Supplementary Table 1 and Methods). These scores were then binarized (1 if the score in a given spot is higher than 1 standard deviation above the mean and 0 otherwise), and we ran pairwise correlation analysis with the Bernoulli model to obtain metabolite groups (Supplementary Table 1 and Methods; **Test statistic 4**). This analysis identified four groups of co-localized metabolites (Fig. 3A) whose respective regions of activity aligned closely with the four regions identified in our zonation based on our pairwise analysis of enzyme co-localization in Figure 2C (Fig. 3B-C). The tumor region (Module 1) was associated with exchange of L-lactate, a well-established hallmark of cancer metabolism [74, 75] (Fig. S5; continuous tissue scores obtained from **Test statistic 8**). This hypothesis was driven by co-expression of lactate transporters from the solute carrier family (*SLC16*) that was largely confined to the tumor region (Fig. 3D; tissue scores obtained from **Test statistic 7**) [76–78]. Additional metabolites enriched in this region included L-glutamate and pyruvate, which are central to energy metabolism in cancer [79, 80], as well as L-arginine, L-proline, and L-ornithine, which are involved in polyamine biosynthesis, a pathway recognized for its role in tumorigenesis, both in cancer cells and (more recently) in other subsets in the TME [81, 82].

**Figure 3.**
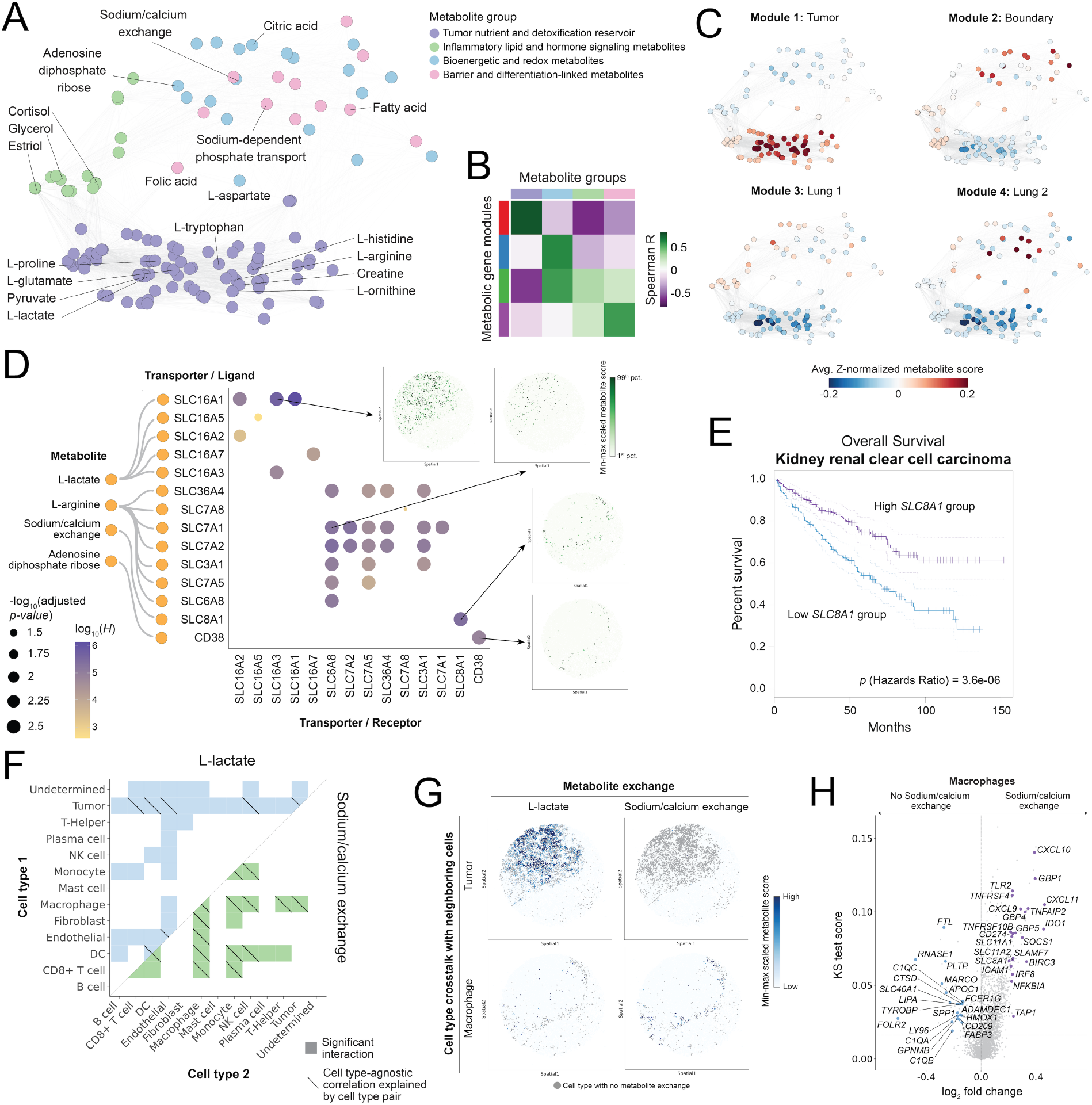
Metabolic crosstalk analysis identifies tumor-associated interactions. (A) Metabolite network colored by metabolite group assignment. Each node represents a metabolite, and edges connect metabolites with a statistically significant (theoretical FDR < 0.05; **Test statistic 4**) pairwise correlation. (B) Cluster map showing the Spearman correlation between the metabolic gene module scores defined in Figure 2 and the metabolite group scores. (C) Metabolite network colored by the average Z-normalized metabolite score in each of the metabolic zones defined in Figure 2. (D) Dot plot showing the statistically significant (empirical FDR < 0.05; **Test statistic 2**) gene pairs for metabolites of interest (selected from **Test statistic 3**). The Harreman statistic and the significance value for the empirical **Test statistic 2** are shown. The spatial plots of some of the relevant gene pairs are shown on the right (**Test statistic 7**). (E) Kaplan-Meier curve of the survival analysis between the *SLC8A1*-high and low groups for kidney renal clear cell carcinoma patients. (F) Heat map with the significant cell type pairs (empirical FDR < 0.01; **Test statistic 6**, empirical null hypothesis 1) for the labeled metabolites. Diagonal lines are added to cell type pairs with a significant contribution to the observed cell type-agnostic autocorrelation of the corresponding metabolite (empirical FDR < 0.05; **Test statistic 6**, empirical null hypothesis 2). (G) Spatial plots of the min-max scaled metabolite scores corresponding to cell types of interest (**Test statistic 10**). Spots labeled as tumor (top) or macrophage (bottom) with no exchange of the corresponding metabolite are colored in gray. (H) Volcano plot showing the macrophage-specific differential expression analysis between macrophage-labeled spots with a positive value for sodium/calcium exchange between macrophages and any other cell type, and the rest of the macrophage-labeled spots. Relevant genes are labeled. KS: Kolmogorov-Smirnov.

Module 2, corresponding to the tumor boundary, was strongly associated with sodium/calcium exchange (NCX) (Fig. 3A, Fig. S5), aligning with recent research on the role of NCX in cancer-associated fibroblasts in modifying migration and consequently reducing cancer cell proliferation [83]. The respective transporter *SLC8A1* (also known as *NCX1*) showed striking spatial specificity for the tumor boundary (Fig. 3D). This gene is expressed on the tumor boundary and is highly prognostic in nasal cancer [84]. Interestingly, while the expression of *NCX1* is not predictive for survival in lung cancer, it correlates positively with survival in RCC, the primary tumor type in this study (Fig. 3E and Fig. S6A). This implies that kidney-derived lung metastases retain characteristics of the primary tumor, including aspects of its metabolic and stromal architecture [85, 86]. More broadly, when looking at the cumulative expression of kidney RCC- and lung-specific genes by defining gene signatures (Methods), we found that the tumor areas (Modules 1 and 2) have a high score of the RCC-specific gene signature, compared to a very low score of the lung-specific signature (Fig. S6B). This supports the notion that metastases originating from kidney RCC may preserve the molecular features of their tissue of origin, even at distant sites, and form a distinct metastatic niche surrounded by RCC-like stroma [86, 87].

### Sodium/calcium exchange via NCX1 may help shape macrophage function in the tumor boundary

To further investigate the cellular context of lactate and sodium/calcium metabolism, we used the next (and finest) level of analysis afforded by Harreman, namely inferring cell-type-specific exchange activity, answering the following question: Is metabolite *m* exchanged by cell types *t* and *u*?, for which the test statistic is computed by summing over all gene pairs (Fig. 3F, Methods, and Supplementary Table 1; **Test statistic 6**, computed by summing over **Test statistic 5**). This analysis identified tumor cells as a primary hub of lactate exchange, particularly in interaction with CD8+ T cells, dendritic cells, and NK cells. Dendritic cells process tumor antigens and play a role in the activation of CD4+ and CD8+ T cells in response to NK-secreted environmental factors. Lactate, though, promotes a tolerogenic phenotype in the TME by leading to an IL-10 production increase, and therefore preventing dendritic cell maturation as well as T cell activation [88]. In contrast, sodium/calcium exchange at the tumor boundary is predominantly associated with macrophages, which communicate with both tumor and immune cells (Fig. 3F). These observations, which are based on spatial data, are supported by a simpler analysis of the single-cell RNA-seq reference, showing the specificity of lactate and sodium/calcium to the respective cell subsets (Fig. S6C).

To better understand the phenotype of macrophages engaging in sodium/calcium exchange at the tumor boundary, we compared their expression profile with that of other macrophages (Fig. 3G-H). For this, we first selected the spots labeled as macrophages (the cell type with the highest z-normalized DestVI-insferred [47] proportion was assigned to each spot; Fig. 3G), and then split them into two groups: (1) the ones with a non-zero and (2) the ones with a zero sodium/calcium exchange score from the metabolite-level and cell-type-specific Harreman analysis (Fig. 3G, Supplementary Table 1 and Methods; **Test statistic 10**). We then performed a differential expression analysis using the macrophage-specific gene expression imputed using DestVI [47] (Fig. 3H). The exchange-active macrophages exhibited a pro-inflammatory program, marked by high levels of the IFN*γ* inducible chemokines *CXCL9, CXCL10*, and *CXCL11* that are specifically recognized by *CXCR3*, and are indicative of an immune-active microenvironment [89, 90]. They also upregulated *SLC11A1* and *SLC11A2*, which are associated with the innate-immune function of macrophages through iron recycling [91, 92], as well as *TAP1*, which supports immune activation [93]. Together, these results are consistent with the roles of sustained calcium flux to enable chemokine production and promote a proinflammatory phenotype in macrophages [94]. More broadly, they support the notion that *NCX1*-mediated control of intracellular calcium balance (through exchange with sodium) impacts macrophage activation [95], and localize it to the promotion of immune activity at the tumor boundary.

### Harreman identifies shared and unique metabolic modules in DSS-induced colitis

We next applied Harreman to study metabolic zonation and exchanges during inflammation in the colon in a Dextran Sulfate Sodium (DSS)-induced colitis mouse model [97]. The dataset consists of two tissue sections, one from each mouse, profiled with 10x Visium (6,175 spots of 55 *µ*m in diameter in total) [96]. The first section was obtained before DSS administration (day 0), and the second one on day 14 after the inflammation peaked and early regeneration began (Fig. 4A). The tissue was rolled from the proximal (outermost part in Fig. 4B) to the distal side (innermost part in Fig. 4B) of the colon, which is commonly known in the field as a Swiss roll. This is a standard histology protocol to fit the entire colon on one slide for posterior sequencing. Histology analysis of the two sections (Fig. 4B) reveals clear effects of damaged tissue, inflammation, and presence of lymphoid patches that likely reflect enhanced recruitment of immune cells [96].

**Figure 4.**
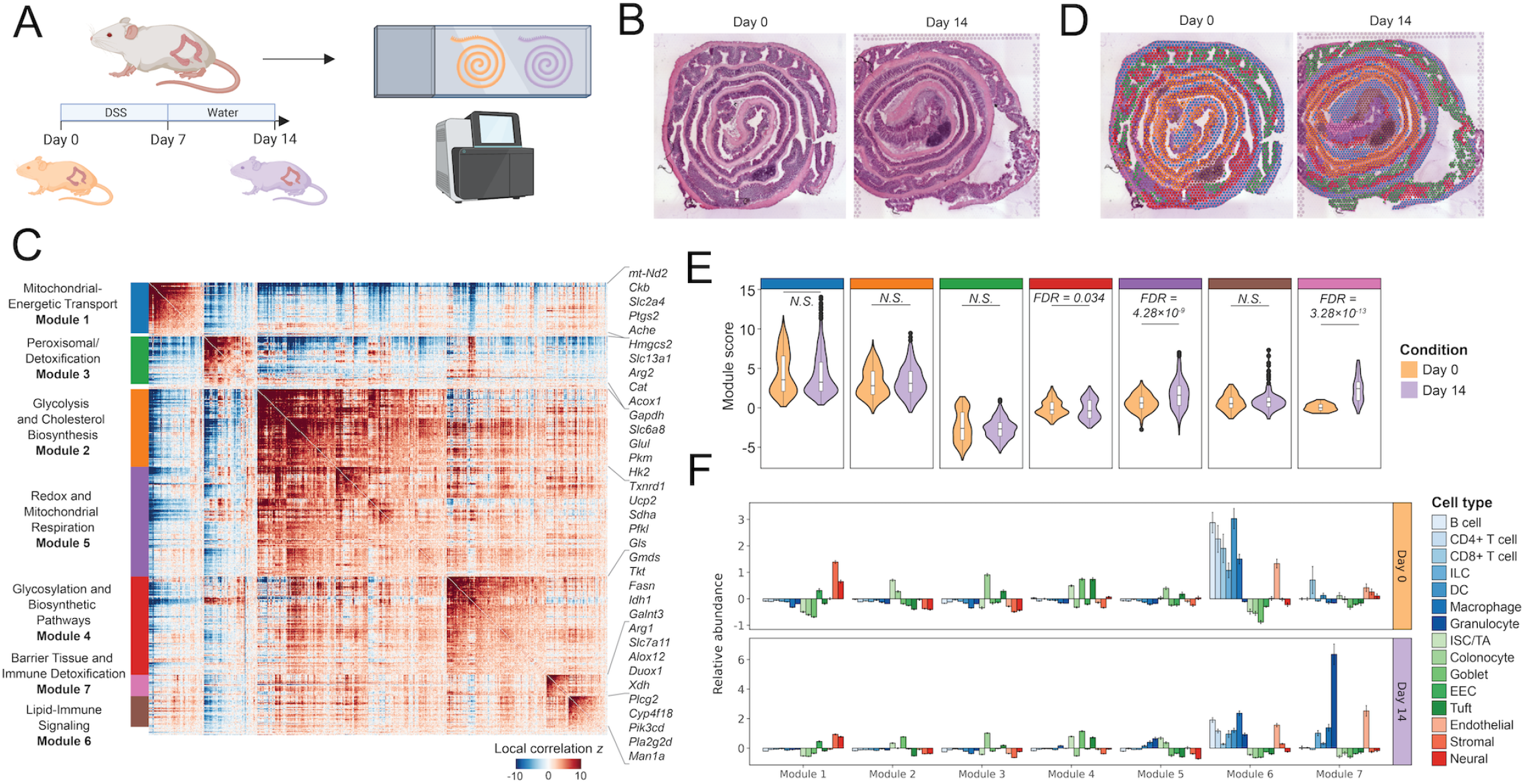
Metabolic reprogramming of the mouse colon in regeneration after inflammation. (A) Schematic representation of the study [96]. Spatial samples were collected on day 0 (WT mouse) and day 14 (regeneration post DSS-induced inflammation) and were profiled with 10x Visium. Figure created using [BioRender](https://www.biorender.com/). (B) Hematoxylin and eosin (H&E) staining of the two samples (day 0 and day 14). (C) Cluster map comprising the 759 metabolic genes with significant autocorrelation (theoretical FDR < 0.01; **Test statistic 1**), grouped into 7 modules (using **Test statistic 2**). (D) Spatial distribution of modules across samples. (E) Violin plots of the module score distribution in the spots with high abundance of the corresponding module (shown in panel D; Methods). Significance computed using the Mann-Whitney U test. P-values were adjusted using the Benjamini-Hochberg approach. N.S.: not significant. (F) Bar plot showing the relative abundance of each cell type across modules for each condition. Error bars represent the standard error of the mean.

The first component of the Harreman pipeline uncovered 7 metabolic modules present across both slides (Fig. 4C-D, Fig. S7A-C, and Methods; **Test statistics 1 and 2**). The colonic Swiss rolls were computationally unrolled to determine the proximal-distal and serosa-luminal axes (Fig. S7D-E and Methods). Because the unrolled coordinates preserve *in vivo* spatial relationships more faithfully than the histological Swiss roll, subsequent spatial analyses were conducted on the unrolled colon coordinates. We confirmed that similar results are obtained when using the original Swiss roll spatial coordinates, suggesting that the spatial neighborhood is generally restricted to those spots present in the same layer (Fig. S8, Methods). We further confirmed that the modules we detected were robust to changes in the number of neighbors considered for each cell as well as the minimum number of genes selected for module computation (Fig. S9, Methods).

The inferred modules showed region-specific (Fig. S7F) and condition-specific (Fig. 4E, Fig. S7C) spatial patterns, characterized by distinct cell type compositions (Fig. 4F) and functional profiles (Fig. 4C). While Module 1 delineates the stromal/neural region (Fig. 4F, Fig. S7C,G), Modules 2 and 3 show mid-distal and proximal colon-specificity, respectively (Fig. 4D, Fig. S7C,F), delineating, together with Module 4, the epithelial/stromal regions of the colon in both health and disease (Fig. 4F). Modules 5, 6, and 7, on the other hand, reflect immune-related processes in the healthy colon (Module 6; Fig. 4D,E) as well as during regeneration after DSS-induced inflammation (Modules 5 and 7; Fig. 4D,E). Supplementary Table 2 provides more information on the functional characteristics and relevance of these modules.

Since Modules 5-7 represent immune- or inflammation-associated modules, with Modules 5 and 7 being mainly present in the distal part of the inflamed condition, we put a special focus on these tissue zones. The metabolic profile of Module 5 suggests a hypermetabolic, oxidative stress-adapted state in the squamous/hyperplastic region of regenerating tissue. For example, *Ucp2* is upregulated in colitis as a mechanism to mitigate reactive oxygen species, and studies in DSS colitis models show that the expression of this gene helps dampen inflammatory damage by limiting oxidative stress [98]. Further, the presence of *Txnrd1*, a key antioxidant enzyme whose induction is a hallmark of cells responding to high oxidative loads [99], indicates activation of antioxidant defenses in the squamous-regenerating epithelium. The inclusion of *Gls, Sdha*, and *Pfkl* reflects high mitochondrial respiration and glycolytic flux, which is common in proliferating or stressed cells that require both ATP and biosynthetic precursors. Indeed, *Gls1* expression increases in “inflammatory hyperplasia” and neoplastic states, indicating greater glutamine utilization when cells are rapidly proliferating under stress [100]. Module 6 was highly correlated with regions rich in immune cells both in the healthy and inflamed conditions, including some metabolic genes predominantly associated with immune activation and survival, such as *Plcg2* and *Pik3cd. Pla2g2d* is an enzyme that acts in lymphoid and inflamed tissues to generate anti-inflammatory lipid mediators to suppress excessive T cell responses and actively resolve inflammation [101]. Likewise, *Cyp4f18* is a cytochrome P450 enzyme expressed in neutrophils that inactivates leukotriene *B*_4_. The degradation of this neutrophil chemoattractant eventually tamps down neutrophil-driven inflammation [102]. Module 7 was highly abundant in the day-14 regenerative squamous areas, which were characterized by abundant granulocytes (neutrophils) and reactive stromal changes (edema in deeper layers). The genes in this module indicate a coordinated response between infiltrating myeloid cells and the stressed epithelium to restore barrier function and detoxify harmful metabolites. For example, *Arg1* is a marker of anti-inflammatory myeloid cells that promotes mucosal healing [103]. In the context of DSS-induced colitis, researchers showed that high levels of this gene favored the accumulation of IL-17A but reduced the expression of IL-17F, eventually alleviating inflammation [104]. *Slc7a11* (also known as *xCT*) encodes the cystine–glutamate antiporter, which is vital for glutathione synthesis in cells under oxidative stress and plays a role in the suppression of ferroptosis in colitis [105]. Further, the module’s inclusion of *Duox1* suggests active production of reactive oxygen species at the mucosal surface [106].

We further characterized the modules by looking at cell type composition (Fig. 4F, Methods). Modules 5-7 were highly correlated with the abundance of immune cells, particularly of granulocytes, reflecting the immune reprogramming that occurs during inflammation and subsequent tissue regeneration. This is in line with literature since in the DSS-induced colitis model, granulocytes, particularly neutrophils, are a key component of the inflammatory response [107, 108].

In summary, Modules 5-7 represent three tissue regions that reflect immune responses. In both conditions, immune cell-rich regions exhibit specialized lipid signaling (Module 6), while, during colitis regeneration, a unique hypermetabolic epithelial state (Module 5), as well as cooperative barrier defense programs with granulocytes (Module 7), emerge.

### Granulocytes exchanging Vitamin A and Lysophosphatidylcholine are key players in gut inflammation and remission phases

We then moved from the enzyme-centric perspective to the metabolite-centric one to uncover metabolites that are being exchanged in a zone-specific manner (Methods and Supplementary Table 1; **Test statistic 3**). For this, we computed metabolite groups (Fig. 5A) using the same pipeline as for the previous dataset and identified five colocalized metabolite exchange groups that were uniquely associated with the two conditions (Fig. 5B, Methods and Supplementary Table 1; **Test statistic 4**). Additionally, the regions of activity of these groups aligned closely to the zones we defined with the enzyme-centric approach (Fig. 5C). In particular, the metabolic module delineating the proximal region of the colon (Module 3) was strongly associated with the group consisting of bile acids, cholesterol derivatives, and TCA cycle intermediates, among others (Fig. 5A,C), possibly reflecting metabolic communication between colonocytes and the environment rather than primary absorption of these metabolites, which occurs mainly in the small intestine [110]. In the mid-distal colonic region defined by Module 2, metabolites mirroring glycolytic reliance, such as ketone bodies, pyruvate, and lactate, as well as microbial metabolites fueling colonocyte energy needs, such as butyrate and acetate [111], were particularly abundant, reflecting the spatial co-expression of their associated transporters (Fig. 5A,C).

**Figure 5.**
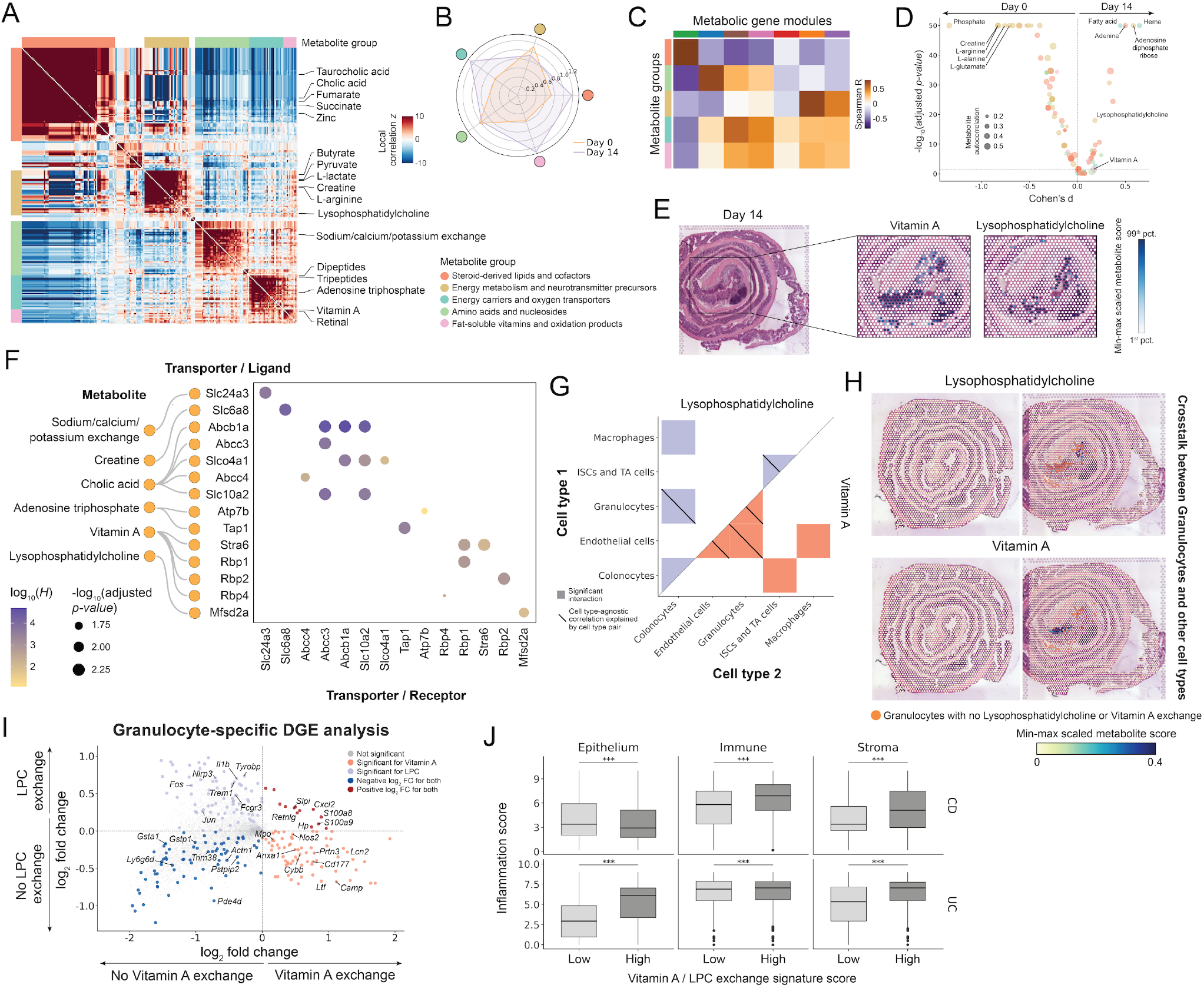
Metabolic crosstalk analysis in the mouse colon reveals region and condition-specific interactions. (A) Cluster map of the pairwise binarized metabolite score correlation with the metabolites grouped into 5 groups (using **Test statistic 4**). (B) Circular plot showing the fold enrichment of metabolite groups in the selected spots (those with a value higher than 1 standard deviation above the mean) for every condition. (C) Cluster map showing the Spearman correlation between the metabolic gene module scores defined in Figure 4 and the metabolite group scores. (D) Volcano plot showing the differential metabolite analysis between day 14 and day 0. Metabolites are colored by the group they belong to, as in (A). Dot size is proportional to the metabolite spatial autocorrelation. Relevant metabolites are labeled. (E) Min-max normalized metabolite scores for vitamin A and lysophosphatidylcholine in a region of interest in the distal part of the regenerating tissue on day 14 (**Test statistic 8**). (F) Dot plot showing the statistically significant (empirical FDR < 0.05; **Test statistic 2**) gene pairs for metabolites of interest (selected from **Test statistic 3**). The Harreman statistic and the significance value for the empirical **Test statistic 2** are shown. (G) Heat map with the significant cell type pairs (empirical FDR < 0.01; **Test statistic 6**, empirical null hypothesis 1) for the labeled metabolites. Diagonal lines are added to cell type pairs with a significant contribution to the observed cell type-agnostic autocorrelation of the corresponding metabolite (empirical FDR < 0.05; **Test statistic 6**, empirical null hypothesis 2). (H) Spatial plots of the min-max scaled vitamin A and lysophosphatidylcholine scores corresponding to granulocytes (**Test statistic 10**). Granulocyte-labeled spots with no exchange of the corresponding metabolite are colored in orange. (I) Scatter plot showing the granulocyte-specific differential expression analysis between granulocyte-labeled spots with a positive value for vitamin A exchange between granulocytes and any other cell type, and the rest of the granulocyte-labeled spots (x-axis), and the same analysis for lysophosphatidylcholine (LPC) (y-axis). Relevant genes are labeled. (J) Box plots showing the inflammation score distribution across disease groups (ulcerative colitis (UC) and Crohn’s disease (CD)) and cellular compartments (epithelium, immune, and stroma) for the bottom and top 20% of the vitamin A/LPC signature score distribution (labeled as “Low” and “High”, respectively) in a human single-cell dataset [109]. *Mann-Whitney U test FDR < 0.05; ** < 0.01; *** < 0.001.

Among the metabolite groups that have an over-representation of spots in the day 14 condition, the activity of the “Fat-soluble vitamins and oxidation products” group (Fig. 5A) was almost completely confined to the granulocyte-rich regenerating region defined mainly by Module 7 (Fig. 5B-C). The metabolites that belong to this group are vitamin A, vitamin D3, retinal, and L-cystine, among others, pointing to barrier repair, induction of a suppressive immune phenotype, and redox protection [112, 113] (Fig. 5A). Interestingly, supplementation of vitamin A in human ulcerative colitis patients has been associated with better clinical response (based on Mayo Clinic score and subscores) and improved mucosal healing compared with the placebo group [114], supporting its observed increase during the regeneration phase (Fig. 5B).

In addition to characterizing day 14-specific metabolite groups, we next investigated which individual metabolites were significantly enriched at day 14 compared to day 0, and which displayed distinct spatial reorganization between these time points (Fig. 5D). Unlike other metabolites that were also significantly overabundant in the day 14 condition, vitamin A and lysophosphatidylcholine were almost absent in the WT tissue and their region of activity was restricted to the distal part of the regenerating colon (Fig. 5E, Fig. S10; continuous tissue scores obtained from **Test statistic 8**), leading us to look deeper into them.

Vitamin A is known to modulate intestinal inflammation by maintaining immune tolerance. Specifically, its active form, retinoic acid, promotes the differentiation of regulatory T cells and IgA-producing B cells, both of which are key players in suppressing excessive inflammation and supporting mucosal immunity [115, 116]. Furthermore, vitamin A contributes to the structural integrity of intestinal epithelial cells, reinforcing the epithelial barrier [117, 118]. Deficiency in vitamin A has been linked to increased intestinal permeability (“leaky gut”), therefore promoting inflammation [112].

In contrast, the role of LPC in gut inflammation is less well understood. However, its transporter, *Mfsd2a* (Fig. 5F), is known to facilitate the uptake of DHA (docosahexaenoic acid) into endothelial cells in the form of LPC–DHA [119]. *Mfsd2a* is essential for maintaining DHA levels in the vasculature, enabling the production of anti-inflammatory lipid mediators such as resolvins, protectins, and maresins [120]. In the context of colitis, a study showed that *Mfsd2a* is essential for maintaining DHA-derived pro-resolving lipid mediator production in the gut vasculature, and that restoring the function of this transporter in endothelial cells promotes resolution of intestinal inflammation in ulcerative colitis and DSS-induced colitis models [121]. Our findings therefore support a potential role for LPC as a key regulator of the resolution phase of gut inflammation.

To explore which cell types may mediate LPC and vitamin A metabolism in the context of inflammation, we performed the last step of the Harreman pipeline to characterize the cell-type-aware communication events by answering the following question: Is metabolite *m* exchanged by cell types *t* and *u*?, for which the test statistic is computed by summing over all gene pairs (Fig. 5G, Methods, and Supplementary Table 1; **Test statistic 6**, computed by summing over **Test statistic 5**). Despite the analysis being performed on all possible cell type pairs, we focused on granulocytes, given their established role in gut immune regulation [122] and their involvement in the exchange of both metabolites (Fig. 5G). Given the overlapping spatial expression of LPC and vitamin A (Fig. 5E), and their association with granulocytes (Fig. 5G), we examined potential co-localization and cell–metabolite interactions. In Fig. 5H, we visually compare granulocytes from day 0 and day 14 (Supplementary Table 1 and Methods; **Test statistic 10**). At day 0, granulocytes were largely absent, whereas by day 14, they were readily detectable and showed distinct regions of co-localization between vitamin A and LPC signals.

To further characterize these granulocyte populations, we performed a differential expression analysis between granulocytes actively exchanging vitamin A and/or LPC and those not involved in exchange, revealing a distinct granulocyte subtype (Fig. 5I). In particular, granulocytes involved in the exchange of both vitamin A and LPC exhibited a repair-oriented inflammatory gene expression profile suggestive of a subset specialized in antimicrobial defense and dampening protease-driven tissue damage (*S100a8/9*), as well as in chemotaxis (*Cxcl2*) and tissue repair and resolution (*Retnlg* and *Hp*). Especially, *S100a8/9* are very strong indicators of inflammation, also used in clinics as biomarkers for IBD activity [123–125]. This is in line with literature that points out a crucial role for neutrophils in the acute phase of colitis inflammation and the initial phase of remission [126]. Compared to the subset exchanging both metabolites, the subpopulation that does not exchange either vitamin A or LPC lacks strong canonical neutrophil granule markers (like *Mpo, S100a8, Prtn3*). However, it does carry a set of genes involved in redox metabolism (*Gsta1, Gstp1*), cytoskeletal remodeling (*Actn1, Pstpip2*), and signal regulation (*Pde4d, Trim38*), suggesting this group could represent a less antimicrobial, more regulatory/metabolic neutrophil state.

Finally, we applied these findings to a human single-cell dataset [109] to uncover if the murine transcriptomic profile corresponding to granulocyte spots with vitamin A and LPC exchange is associated with high inflammation conditions. We performed this analysis separately for each cellular compartment (epithelium, immune, and stroma) and IBD disease (ulcerative colitis (UC) and Crohn’s disease (CD)), and almost all cases showed an increase in inflammation score for the population with the highest vitamin A/LPC exchange signature score compared to the population with the lowest score (Fig. 5J, Methods). This means that the inflammatory behavior is still imprinted in the day 14 tissue, despite it being in the regenerative stage.

## Discussion

We introduce Harreman, a computational method that infers metabolic exchange in spatial transcriptomics data. Harreman builds on the Hotspot algorithm, previously developed by our group [35]. We present a layered way to analyze metabolic spatial correlation, by first inferring metabolic zones by using a metabolic enzyme-centric perspective and then studying the metabolite exchanges in the predefined tissue zones. We adapted the algorithm to perform cell-type-aware communication inference by evaluating the pairwise spatial correlation of gene pairs between spatially proximal cells. To run Harreman effectively and scale it to data sizes characteristic of state-of-the-art spatial transcriptomics technologies such as Slide-seq [45], we made it PyTorch-friendly, which led us to the subsequent optimization of the Hotspot algorithm using the same infrastructure.

We applied Harreman in the context of RCC-derived lung metastasis and DSS-induced inflammation and found important spatially- and cell-type-dependent metabolic processes. In particular, sodium/calcium exchange through the *SLC8A1* transporter was shown to shape macrophage phenotype in the tumor boundary by increasing its inflammatory behavior. In the context of colitis, our analysis revealed that metabolites such as vitamin A and lysophosphatidylcholine were highly abundant in the regenerating tissue following DSS-induced barrier damage, especially in the granulocyte-infiltrating areas, elucidating metabolic crosstalk events that might play a role in tissue regeneration after inflammation.

Harreman introduces a cell-type-dependent (as well as an independent) statistic that is based on spatial autocorrelation and therefore accounts for data sparsity commonly found in low-sensitivity spatial technologies. To support common spot-based spatial transcriptomics technologies, deconvolved results using a reference single-cell dataset can also be fed into the algorithm to gain additional resolution by imputing cell-type-specific gene expression data in every spot. Further, metabolite exchange-related databases are currently lacking, and in this work, we addressed this limitation by building HarremanDB, a metabolite transporter database consisting of manually curated and publicly available metabolite-transporter relationships [29, 28, 36–42]. Pairwise transporter co-expression and their spatial localization can be leveraged to identify metabolite-exchanging cells or spots at an individual level, with the eventual possibility to study the transcriptional profile of those cells by performing a differential expression analysis.

Metabolic zonation performed with Harreman provided a deeper biological understanding of both general and specific metabolic processes. This was particularly striking in the comparison between healthy and DSS-treated colon, where we identified metabolic modules such as Modules 1, 2, and 3 that were not only shared between the two conditions but also were spatially constrained to the stromal, mid-distal, and proximal regions, respectively. In contrast, Modules 5 and 7 were specific to the inflamed gut. Notably, the metabolic processes in Module 7, being confined to a very limited spatial area, could easily be overlooked or obscured when analyzing the data as a whole. The identification of known metabolites involved in healthy vs. inflamed gut, such as vitamin A, increases our confidence in the reliability of Harreman and its potential for discovering novel findings. It was precisely the stratification of metabolic zones that enabled us to detect LPC as a key metabolite in both inflammation and remission.

The biological value of Harreman also lies in its ability to associate metabolic processes with the cell types directly responsible for them, offering a more detailed view of tissue organization. This is especially relevant in contexts such as tumors and metastases, where tumor–stroma interactions play a crucial role in disease aggressiveness and progression. Identifying which cells are responsible for specific metabolic processes that support tumor growth opens new avenues for targeted therapies. In our dataset, we were able to confirm that RCC-derived lung metastases retained the primary tumor’s metabolic signature, suggesting a metabolic continuity that may be critical for metastatic colonization and offering a potential target for intervention. In particular, we identified *SLC8A1* (also known as *NCX1*), a sodium/calcium exchange transporter known to correlate with better prognosis in RCC. Further investigation into the role of this transporter in RCC-derived metastases could reveal whether it represents a protective factor, a therapeutic target, or a biomarker for disease progression.

Limitations of Harreman include the assumption that transporter expression reflects metabolite presence and its import from or export to the surrounding environment, without considering the intracellular metabolic pathways and post-translational modifications that are required for that to happen [127, 128]. The algorithm is currently dependent on the curated metabolite transporter (as well as the CellChat-derived [43] ligand-receptor) database from which the interactions are taken, which may represent a subset of the whole set of possible transporters that exchange a given metabolite. The actual version of the database is also agnostic of the difference between import and export reactions, as all transporters are considered to be able to do both. However, further refinement would be helpful to improve the algorithm’s inference capabilities.

We made the software available at https://github.com/YosefLab/Harreman with its accompanying tutorials (https://harreman.readthedocs.io/) so that it can be easily accessible for the community.

## Methods

### Optimization of the Hotspot algorithm

Due to the exponential scaling of the single-cell profiling technologies [129], there is an increasing number of cells that are being sequenced nowadays. Therefore, computational resources have to be adapted to the required needs, and the algorithms have to be designed in such a way that they are able to deal with the increasing number of cells. This motivation led to the optimization of the Hotspot algorithm [35] by using PyTorch.

The equations that have been used to optimally run Hotspot are described in the “Representing the equations as matrix multiplications” section (*Test statistics 1 and 2*).

We compared the optimized version (implemented inside Harreman) with the original Hotspot algorithm (Fig. S1). For this, we used a set of 1,547 CD4+ T cells filtered from a sample of human peripheral blood mononuclear cells (PBMCs) (made available by 10x Genomics) as well as the Slide-seq dataset [45] that were used in the original publication [35].

In the spatial Slide-seq dataset, the *Puck_180819_12* sample was considered (32,701 spots). To create the neighborhood graph and compute gene autocorrelations, Harreman and Hotspot were run with the following parameters: *n_neighbors=300, weighted_graph=False*, and *model=‘bernoulli’*. Further, the *approx_neighbors=False* parameter was also used in Hotspot to make sure the kNN graphs generated by both methods are identical. Local correlations were computed for all the significantly autocorrelated genes (876 genes; FDR < 0.05). To create gene modules, the *min_gene_threshold=20, core_only=False*, and *fdr_threshold=0*.*05* parameters were used, ending up with a total of 6 modules (Fig. S1A-C).

In the single-cell PBMC dataset, principal component analysis (PCA) with 10 components was performed to compute the transcriptomic latent space. To create the neighborhood graph and compute gene autocorrelations, Harreman and Hotspot were run with the following parameters: *n_neighbors=30, weighted_graph=False*, and *model=‘danb’*. Further, the *approx_neighbors=False* parameter was also used in Hotspot to make sure the kNN graphs generated by both methods are identical. Local correlations were computed for the top (highest Z value) 500 significantly autocorrelated genes (FDR < 0.05). To create gene modules, the *min_gene_threshold=15, core_only=True*, and *fdr_threshold=0*.*05* parameters were used, ending up with a total of 13 modules (Fig. S1D-F).

Apart from the mentioned datasets, two smaller versions of the Slide-seq dataset (21,667 and 11,902 spots) were also used to run Harreman and Hotspot on them and evaluate the runtime per CPU (or GPU in the case of using the PyTorch-based Harreman version; Fig. S1G). For this, the same parameters as the ones described in the previous paragraph were used. However, the top (highest Z value) 500 significantly autocorrelated genes (FDR < 0.05) were used in all datasets to avoid differences in runtime due to a different number of genes to evaluate pairwise correlations on. We then evaluated the effect of the number of autocorrelated genes on the overall runtime by running Hotspot and Harreman on the spatial Slide-seq dataset on a varying set of genes (Fig. S1H). Note that this procedure only affects the computation of the pairwise correlations, since gene autocorrelation is assessed on the whole set of genes. For this, we selected the top (highest Z value) *n* significantly autocorrelated genes (FDR < 0.05), where we used different values for *n*: 200, 400, 600, and 800. The parameters used were the same as the ones specified in the previous paragraph. To make sure we computed the runtime per CPU, the *jobs=1* parameter was used in Hotspot in both the *compute_autocorrelations* and the *compute_local_correlations* functions in all datasets.

To verify that the theoretical null hypothesis of the original Hotspot algorithm is correlated with the empirical one in the optimized version, the gene autocorrelation and pairwise correlation results were computed using both approaches. The p-values obtained from both approaches were compared against each other (Fig. S1B,E). For this, both the single-cell PBMC dataset and the spatial Slide-seq dataset were used.

### List of metabolic genes for tissue zonation

To perform metabolic zonation of the tissue, a list of metabolic genes has been added. To obtain the list of all metabolic genes for both humans and mice, the Compass tool (version 1.0.0) [130] was used. Compass infers the metabolic state of each cell by solving a linear programming problem, utilizing both transcriptomic data and the metabolic network to output a matrix of reaction scores for each metabolic reaction in each cell. Therefore, the list of metabolic genes that we obtain represents the set of metabolic enzymes that participate in at least one metabolic reaction. For this, the *–list-genes* argument along with the *–species* argument with either the *mus_musculus* or *homo_sapiens* option was used for mouse and human, respectively. Eventually, after manual filtering of unnamed metabolic enzyme genes as well as conversion of the list of mouse genes to the correct nomenclature, 1337 and 1323 metabolic genes have been considered for human and mouse, respectively. For gene name conversion, the *useEnsembl* and *getBM* functions from the *biomaRt* R package were used with the following parameters: *biomart = “ensembl”* and *dataset = “hsapiens_gene_ensembl”* for the *useEnsembl* function, and *filters = “ensembl_gene_id”, attributes = c(“ensembl_gene_id”, “external_gene_name”, “mmusculus_homolog_ensembl_gene”, “mmusculus_homolog_associated_gene_name”)* for the *getBM* function. Further, the human gene list with gene SYMBOL IDs downloaded from Ensembl (GRCh38.p14) [131] was used.

### Metabolic interaction database (HarremanDB)

To assess cellular crosstalk on a reduced set of gene pairs, a gene pair database based on previous knowledge is used as input. For this, two different databases have been created for human and mouse. On the one hand, we built a metabolite transporter database by collecting information from different sources. These are HMDB [36], MEBOCOST [29], TransportBD [37], MetallinksDB [28], RESOLUTE [38], and different scientific articles with manually curated metabolite-transporter information [39–42].

When it comes to HMDB, out of the total number of 217,920 metabolites, those associated with at least one transporter gene have been initially considered (16,007 metabolites). To only focus on extracellular metabolites, those present in the extracellular space and either in the blood or the cerebrospinal fluid have been selected (997 metabolites, both for humans and mice). To generalize the database to the mouse, the mouse genes equivalent to the human genes present in the original database have been considered. However, to get rid of unknown metabolite names and lowly relevant substrates, HMDB metabolites have been manually curated. From the MEBOCOST database, genes annotated as transporters and their respective metabolites have been considered, which were initially curated from scientific articles as well as the Recon2 resource [132] by using the Compass tool [130]. The authors retained those metabolites in “extracellular space”, “blood”, or “cerebrospinal fluid” and were named “extracellular metabolites”. 106 metabolites have been considered for humans and mice. Additionally, we collected metabolite-transporter information from four different studies [39–42], two of them being solute carrier (SLC) transporter reviews [39, 40], to also contain highly reliable annotations. Metabolite nomenclature was slightly modified to match the labels from the other databases. To further extend the human version of HarremanDB, transporter information from TransportDB has been used (downloaded from https://www.membranetransport.org/transportDB2/index.html). To link the transporter substrates with their corresponding genes, we integrated the “TransportDB2.0_translated.tsv” file from the MetalinksDB [28] platform (downloaded from https://github.com/dbdimitrov/metalinks/tree/main/data), as well as the information relating UniProt IDs [133] with gene SYMBOL IDs (the latter downloaded from Ensembl (GRCh38.p14) [131]) using the “org.Hs.eg.db” [134], “EnsDb.Hsapiens.v79” [135], and “biomaRT” [136] R packages. After that, to avoid redundancy, we considered those transporters that were not present in the other datasets that we collected, and to further ensure the reliability of the results, we only considered the transporters that are present in the plasma membrane [137–142], and therefore involved in extracellular transport. Eventually, we manually curated the metabolite annotation, and we integrated the TransportDB metabolite information with the rest of the database. Despite the TransportDB gene names being for humans, we used the equivalent mouse names and also included this database in the mouse version. The last resource that we considered was RESOLUTE (https://re-solute.eu/), which contains a database of 446 human SLCs. To integrate it with HarremanDB, we considered those genes that were not present in the other resources and are located in the plasma membrane. This last information, as well as a coarse identification of the substrates that are carried by each transporter, is present in the RESOLUTE knowledge base (https://re-solute.eu/knowledgebase). The substrate information was further curated manually to make sure the names matched what was provided by the rest of the HarremanDB resources. Similar to what we did with TransportDB, we also generalized the RESOLUTE transporter information from human to mouse.

After manual curation to make sure the metabolite names are unique and interpretable, all these different datasets have been concatenated to build HarremanDB. Two versions of the database have been created, one only containing extracellular metabolites and transporters (241 human and 240 mouse metabolites; used by default), and another one also considering those HMDB metabolites not limited to the extracellular space (1,038 human and 1,037 mouse metabolites).

This database contains information regarding metabolites and the corresponding genes associated with each metabolite. Therefore, gene pairs are created by considering each metabolite individually and making sure that these pairs are unique across the entire dataset. If they are not, this could be interpreted as a given gene pair carrying more than one metabolite.

On the other hand, an LR database has been collected from CellChatDB (version 2.1.2) [43], consisting of 3,379 and 3,233 LR pairs for mouse and human, respectively. Each LR pair belongs to a given pathway, and this information is used to group LR pairs in the same way transporter pairs are linked to a given metabolite. CellChatDB contains 293 and 290 pathways for mouse and human, respectively.

### The Harreman algorithm for layered analysis of metabolic zonation and crosstalk in spatial data

Harreman provides a series of formulas to perform spatial correlation and characterize the metabolic state of tissue using spatial transcriptomics. At the coarsest level, Harreman partitions the tissue into modules of different metabolic functions based on enzyme co-expression. At the following stage, Harreman formulates hypotheses about which metabolites are exchanged across the tissue or within each spatial zone. This is achieved by computing an aggregate spatial correlation that takes into account all gene pairs associated with the import or export of a given metabolite. Spatially co-localized metabolites can also be grouped together. Moving to a finer resolution, Harreman can also infer which specific cell subsets participate in the exchange of distinct metabolic activities inside each zone. In this case, the aggregate spatial correlation is calculated by considering cell-type–restricted expression patterns of import–export gene pairs. Beyond these central functionalities, which are the main focus of this work, Harreman also provides a range of aggregate spatial correlation statistics designed to capture diverse interaction scenarios. In the following, we go over all the layers of analysis as depicted in Supplementary Table 1.

In all the equations below, for proteins composed of multiple subunits, we compute either an algebraic or a geometric mean of the expression values of the corresponding genes as done in SpatialDM [34]:

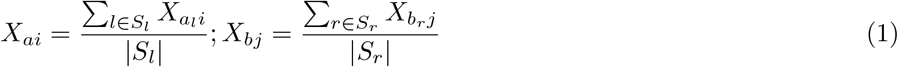

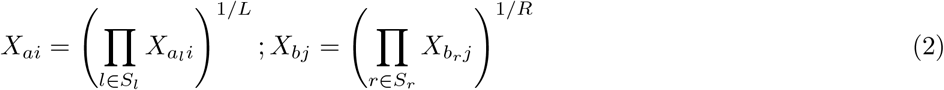

where *l* is a subunit for the protein encoded by gene *a* that belongs to the set of subunits *S*_*L*_, and *r* is a subunit for the protein encoded by gene *b* that belongs to the set of subunits *S*_*R*_. |*S*_*l*_| and |*S*_*r*_| denote the number of subunits for proteins encoded by genes *a* and *b*, respectively.

#### Test statistic 1: Is gene *a* spatially autocorrelated?

When performing tissue zonation, having a limited set of informative genes is useful to constrain the analysis to spatially autocorrelated genes. The set of genes used for the equation depends on whether tissue zonation or cellular crosstalk (explained in **Test statistic 2**) is inferred. In the former case, the set of metabolic genes used by the Compass algorithm [130] is considered (explained in the “List of metabolic genes for tissue zonation” subsection above), while for the latter case, we use the transporters and ligand-receptor (LR) pairs present in HarremanDB and CellChatDB [43], respectively (explained in the “Metabolic interaction database (HarremanDB)” subsection above). Additionally, this statistic is also used when inferring cell-type-agnostic crosstalk (explained in **Test statistic 2**) but for the case of gene *a = b*. The equation used is equivalent to the one defined in the Hotspot algorithm [35]:

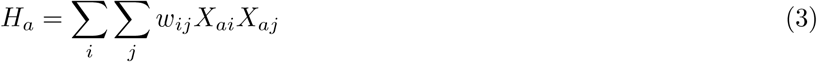

where *a* is a gene expressed by cells *i* and *j*, and *X* refers to the gene expression matrix of dimension genes x cells.

The weight *w*_*ij*_ represents communication strength between neighboring cells. It is positive if *i* and *j* are neighbors and 0 otherwise, and the lower the distance in the similarity graph, meaning a more similar neighbor, the higher the value. To calculate them, a Gaussian kernel is used, which is defined by the following equation:

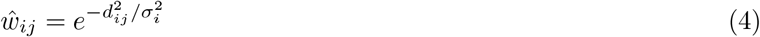

where *d*_*ij*_ corresponds to the distance between two neighboring cells and *σ*_*i*_ refers to the selected bandwidth for cell *i*, which by default is set to *K/*3, where *K* represents the number of chosen neighbors in the K-nearest-neighbors (kNN) graph or the number of neighbors at a distance smaller than *d* micrometers, with the most appropriate values being between 50 and 200 micrometers to focus on local neighborhoods. An unweighted option is also available, where the value is 1 if two cells are neighbors and 0 otherwise.

Weights are also normalized across cells to sum 1 for each cell (this is only done in the weighted case).

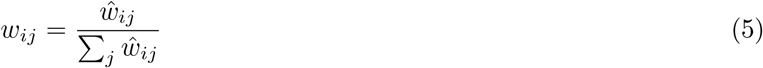

Further, in cases where the dataset contains the spatial positions of two or more samples, to avoid connections between cells or spots that belong to different samples, a weight of 0 will be assigned if two given cells or spots belong to different samples.

To test significance and evaluate expectations for *H*, that is, to assess if the obtained value is extreme compared to what we would expect by chance, a null model is needed. A parametric and a non-parametric approach have been implemented, the former having already been introduced in the Hotspot manuscript [35]. The theoretical null assumes that expression values are drawn independently from some underlying distribution for which we can compute *E* [*X*_*ai*_] and 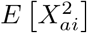 for each cell. The null hypothesis here would be that the expression of gene *a* is independently and identically distributed across cells, that is, that there is no spatial autocorrelation.

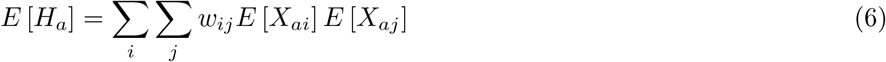

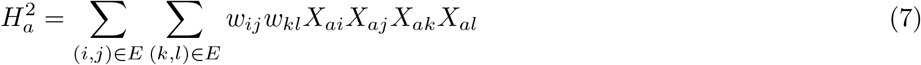

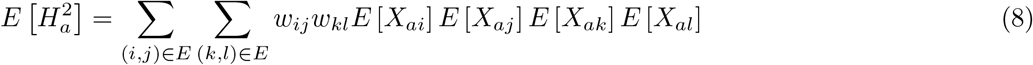

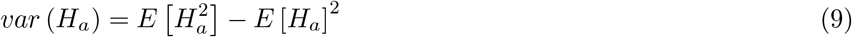

The expression values are standardized before computing the autocorrelation value *H*. For this, different statistical models can be used, where the negative binomial distribution is generally used to model single-cell and spatial data. However, in cases where the counts are very sparse, the Bernoulli distribution might be a better option. Moreover, the normal distribution has also been implemented to model other types of data that don’t follow either of the previously mentioned models [35]. The expression count standardization is performed as follows:

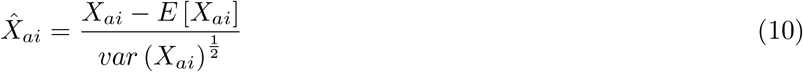

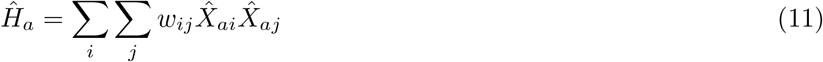

Computing the null model is made simpler as the expectation of *H* is 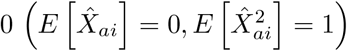 :

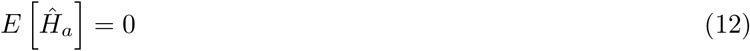

Therefore, the second moment of *H* is as follows:

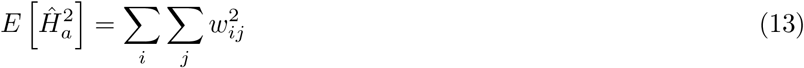

To assess communication significance, *H*_*a*_ is converted into a Z-score using the equation below, and a significance value is obtained for every gene pair by comparing the Z values to the normal distribution:

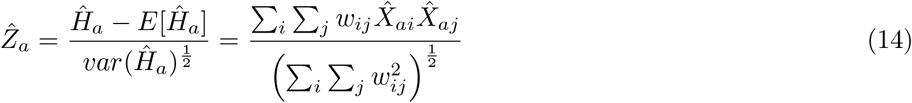

Lastly, p-values are adjusted using the Benjamini-Hochberg procedure [143].

In the empirical test setting, either raw gene counts, log-normalized values, standardized gene expression values, or counts normalized in any other desired way can be used. Cell IDs from the counts matrix are shuffled *M* times (*M* = 1, 000 by default), and the correlation values for each gene according to Eq. 3 are computed in each iteration. Then, Eq. 15 is used to calculate the p-value, where *x* represents the number of permuted *H*_*a*_ values that are higher than the observed one and *M* is the total number of permutations.

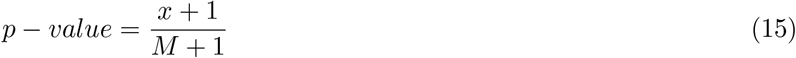

Finally, p-values are adjusted using the Benjamini-Hochberg method [143].

#### Test statistic 2: Are genes *a* and *b* spatially co-localized (or interacting with each other)?

Once metabolic genes with a relevant spatial gene expression pattern have been selected, pairwise correlation can be computed to eventually group genes into modules and define tissue zones. Here we made use of the second formula of the Hotspot algorithm [35], which is defined as follows:

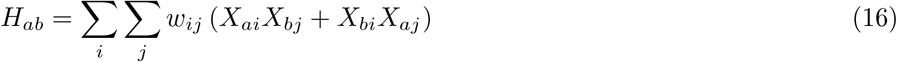

where *a* and *b* are two different genes expressed by cells *i* and *j*, respectively, and *X* refers to the gene expression matrix of dimension genes x cells.

The weight *w*_*ij*_ represents communication strength between neighboring cells, and it is defined in the same way as in **Test statistic 1**.

This statistic is also relevant to address the question of which interactions happen in the defined tissue regions. To infer metabolite exchange (or ligand-receptor interaction) events present in the tissue without adding the cell type constraint, a cell-type-agnostic approach has also been implemented. This analysis, despite not making use of any novel formulas, is based on gene pairs defined in HarremanDB and/or CellChatDB [43], therefore restricting the communication analysis to already defined gene pairs that are known to interact with each other. As gene pairs can be made up of either different or the same genes, the formula needs to be adapted to each case. For the former case, the pairwise correlation formula is used (Eq. 16), while for the latter, that is, if *a* = *b*, the spatial autocorrelation formula defined in **Test statistic 1** is used (Eq. 3).

For significance testing, both an empirical and a theoretical test have been implemented. The latter corresponds to the already existing theoretical test introduced in the Hotspot method [35]. In this case, instead of considering a null model that assumes the expression values of genes *a* and *b* are independent, which significantly underestimates the variance of *H*_*ab*_ if at least one gene has high autocorrelation (which is required to select these genes), a conditionally independent null hypothesis is tested. Here, we test how extreme *H*_*ab*_ is compared with independent values of gene *b* given the observed value of gene *a* (*P* (*H*_*ab*_ *a*)), and vice versa (*P* (*H*_*ab*_ | *b*)). Eventually, we conservatively retain the least-significant result. After going through equations 6-10 adapted for Eq. 16, and conditioning on gene *a*, the second moment of *H* is expressed as follows:

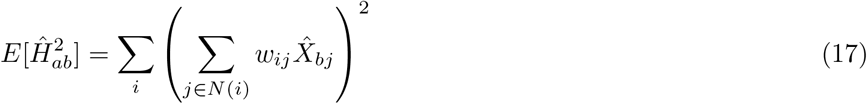

To assess communication significance, the statistic is converted into a Z-score using the equation below, and a significance value is obtained for every gene by comparing the Z values to the normal distribution:

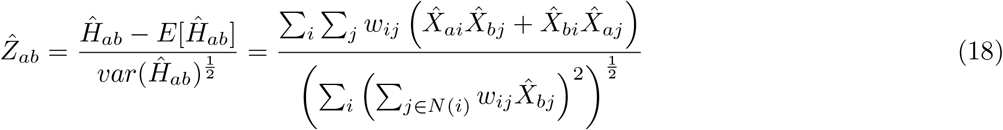

P-values are then adjusted using the Benjamini-Hochberg procedure [143].

In the empirical test setting, cell IDs in the counts matrix corresponding to gene *a* (or *b*) are shuffled *M* times (*M* = 1, 000 by default). Then, the correlation values for each gene according to Eq. 16 are computed in each iteration, and Eq. 15 is used to calculate the p-value, where *x* represents the number of permuted *H* values that are higher than the observed one and *M* is the total number of permutations. Finally, similar to the parametric test, the most conservative p-values are considered. These p-values are then adjusted using the Benjamini-Hochberg method [143].

#### Test statistic 3: Is metabolite *m* spatially autocorrelated?

Apart from getting interaction event information at the gene pair level, those results can be integrated to study one additional layer of intracellular interactions. As each gene pair used to compute *H* is associated with a given metabolite (HarremanDB) or ligand-receptor (LR) pathway (CellChatDB [43]), the above formulas can be summed over all gene pairs that exchange a given metabolite (or interact through a given LR pathway):

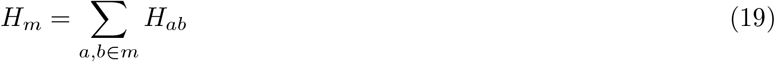

where *m* is a given metabolite that is being exchanged by genes *a* and *b* (or *a* = *b*). In this setting, we would be assessing the spatial autocorrelation of metabolite *m*.

To test for statistical significance using the theoretical test, the same procedure is applied on the second moments of *H*, giving rise to the equation below:

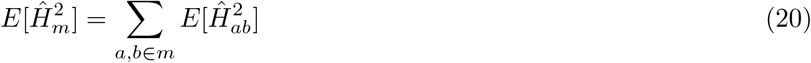

Eventually, Z-scores can be computed using the following equation:

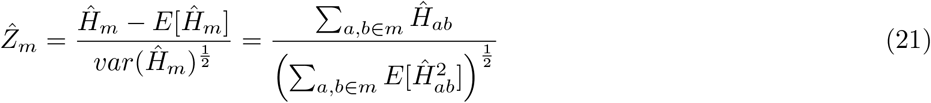

and p-values are obtained for every metabolite by comparing the Z values to the normal distribution. P-values are then adjusted using the Benjamini-Hochberg approach [143].

For significance testing using the empirical test, Eq. 19 is computed in every shuffling iteration. This is done separately for genes *a* and *b* because, when *a* and *b* are different, the cell IDs corresponding to the counts matrices of both gene *a* and *b* are shuffled, giving rise to *H*_*m*_(*a*) and *H*_*m*_(*b*). However, if gene *a* = *b*, then the same output is considered for both *H*_*m*_(*a*) and *H*_*m*_(*b*). Then, two sets of p-values are obtained through Eq. 15, and eventually, the most conservative p-values are selected. P-values are also adjusted using the Benjamini-Hochberg procedure [143].

#### Test statistic 4: Are metabolites *m*_*1*_ and *m*_*2*_ spatially co-localized?

Once we have identified which metabolites show a spatially informed pattern, we can go one step further and group metabolites with similar spatial abundance. For this, a modified version of Eq. 16 defined in **Test statistic 2** is used, which takes as input binarized metabolite scores per cell/spot instead of gene expression data:

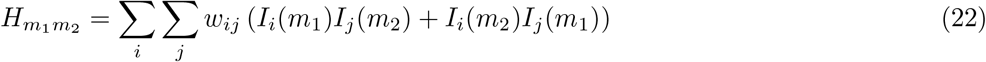

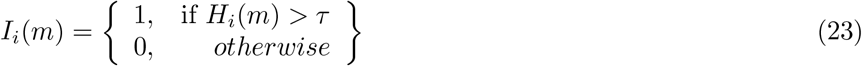

where *τ* is defined as 1 standard deviation above the mean, and *H*_*i*_(*m*) is defined in **Test statistic 8** (Eq. 31).

Similarly to the previous test statistics, the binarized metabolite scores are standardized before computing the pairwise correlation value 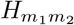. For this, the Bernoulli distribution is used, as it is the best option to model the data. For significance testing, the theoretical and empirical tests implemented in **Test statistic 2** have been adapted to this statistic. In the former case, instead of considering a null model that assumes the binarized scores of metabolites *m*_1_ and *m*_2_ are independent, which significantly underestimates the variance of 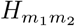 if at least one metabolite has high autocorrelation (which is required to select these metabolites), a conditionally independent null hypothesis is tested. Here, we test how extreme 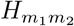 is compared with independent values of metabolite *m*_2_ given the observed value of metabolite 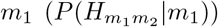, and vice versa 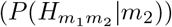. Eventually, we conservatively retain the least-significant result. After going through equations 6-10 adapted for Eq. 22, and conditioning on metabolite *m*_1_, the second moment of *H* is expressed as follows:

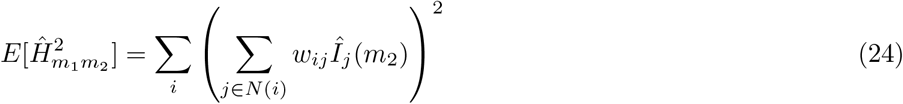

To assess communication significance, the statistic is converted into a Z-score using the equation below, and a significance value is obtained for every gene by comparing the Z values to the normal distribution:

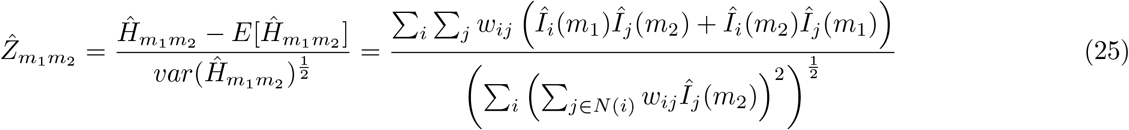

P-values are then adjusted using the Benjamini-Hochberg procedure [143].

In the empirical test setting, cell IDs in the counts matrix corresponding to metabolite *m*_1_ (or *m*_2_) are shuffled *M* times (*M* = 1, 000 by default). Then, the correlation values for each metabolite according to Eq. 22 are computed in each iteration, and Eq. 15 is used to calculate the p-value, where *x* represents the number of permuted *H* values that are higher than the observed one and *M* is the total number of permutations. Finally, similar to the parametric test, the most conservative p-values are considered. These p-values are then adjusted using the Benjamini-Hochberg method [143].

#### Test statistic 5: Do genes *a* and *b* interact when expressed by cell types *t* and *u*, respectively?

To go further and identify the cell types that exchange the most relevant metabolites (or ligand-receptor pathways) in the metabolic regions of interest, a cell type-aware approach has also been developed. The mathematical representation of this test statistic, which is used to quantify the communication strength between a given pair of cell types, is as follows:

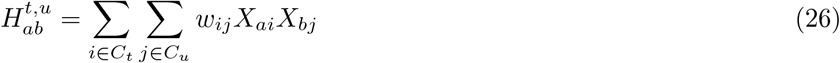

where *i* and *j*, in addition to being two different cells, belong to different cell types *t* and *u*, respectively. *C*_*t*_ and *C*_*u*_ refer to the set of cells that belong to cell types *t* and *u*, respectively. Further, *a* and *b* are two different genes expressed by cells *i* and *j*, respectively, and *X* refers to the gene expression matrix of dimension genes x cells. Further, the expression of the same gene (*a* = *b*) between two different cells must also be considered, i.e., when inferring metabolic crosstalk between cells that express the same transporter.

The weight *w*_*ij*_ represents communication strength between neighboring cells, and it is defined in the same way as in **Test statistic 1**.

Weights are assigned using a spatial proximity graph, such that *w*_*ij*_ is only non-zero if cells *i* and *j* are neighbors and there are no self-edges. This last statement is not true when dealing with deconvolved spot-based spatial data, where interactions between different cell types present in the same spot could be considered. In that case, though, each cell type inferred in each spot using spatial deconvolution methods is treated as a separate node in the graph, where instead of assigning a distance equal to 0, the assigned distance between cell types within the same spot is 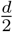, with *d* being the spot diameter. For this, DestVI [47] or cell2location [144] can be used to estimate the cell-type abundance in each spot as well as the cell-type-specific gene expression values. As a result, the double summation can be re-expressed as a sum over edges *E*, which results in the following sparse graph:

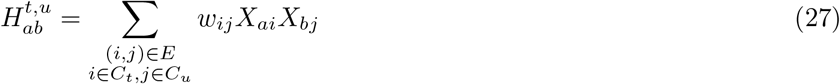

To test significance and evaluate expectations for *H*, a null model is needed. For this, an empirical test has been implemented, where the shuffling procedure varies for each one of the 3 different null models:

1. Given the spatial co-localization of cell types *t* and *u*, which gene pairs are significantly co-expressed by cell types *t* and *u*, respectively? The null hypothesis is as follows: the observed co-expression of gene pair (*a, b*) across cell types *t* and *u* is no stronger than expected by chance, given the spatial co-localization of cell types *t* and *u*. Therefore, gene pair expression counts within their respective cell types are shuffled.
2. Given the spatial autocorrelation of a given gene pair (*a, b*) regardless of cell type, which cell types explain the observed co-localization? The null hypothesis is as follows: the observed co-expression of genes *a* and *b* is not enriched in any specific cell type pair, that is, it is random with respect to which cell types express them. In this case, cell type labels are shuffled.
3. Given a fixed cell type (e.g., stem cells), we test interactions with other cell types. The null hypothesis is as follows: the observed spatial co-expression between gene *a* in a cell type of interest and gene *b* in another cell type *u* is no stronger than expected if gene *b*’s expression were random in cell type *u*. Here, we fix the expression of gene *a* in cell type *t* and shuffle the expression of gene *b* in cell type *u*.

Then, the correlation values for each cell type pair and gene pair according to Eq. 27 are computed in each iteration, and Eq. 15 is used to calculate the p-value. P-values are finally adjusted using the Benjamini-Hochberg approach [143].

#### Test statistic 6: Is metabolite *m* exchanged by cell types *t* and *u* ?

Gene pairs can also be linked to metabolites by using the HarremanDB database (or to ligand-receptor pathways by using CellChatDB [43]). Therefore, as Eq. 26 can be summed up for all gene pairs that exchange a given metabolite, we obtain the equation below:

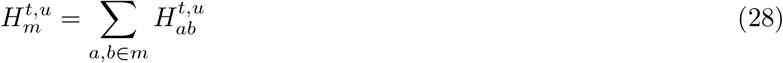

where *m* is a given metabolite that is being exchanged by genes *a* and *b*. In this setting, we would be assessing if metabolite *m* is exchanged by cell types *t* and *u*.

Significance testing in this case also depends on which of the three possible null models defined in **Test statistic 5** we are testing and, therefore, the shuffling strategy used. The null hypotheses in this case are defined as follows:

1. The observed co-expression of all gene pairs associated with metabolite *m* across cell types *t* and *u* is no stronger than expected by chance, given the spatial co-localization of cell types *t* and *u*.
2. The observed co-expression of metabolite *m*’s associated genes is not enriched in any specific cell type pair, that is, it is random with respect to which cell types express those genes.
3. The observed spatial co-expression between the expression of metabolite *m*’s genes in a cell type of interest and the expression of those genes in another cell type *u* is no stronger than expected if gene expression in cell type *u* were random.

However, irrespective of the null hypothesis we want to test, Eq. 28 is computed in each iteration. Eventually, p-values are computed as in Eq. 15 and adjusted using the Benjamini-Hochberg approach [143].

#### Test statistic 7: Do genes *a* and *b* interact when *a* is expressed by cell *i* and *b* by spatially nearby cells?

The results obtained using the cell-type-agnostic approach described in **Test statistics 1 and 2** can be visualized in space in a way such that each cell/spot is assigned a score aggregating the communication with neighboring cells/spots through a given gene pair (*a, b*):

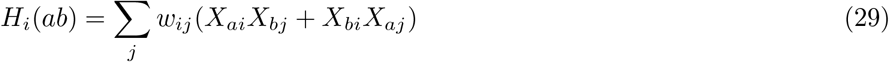

If *a* = *b*:

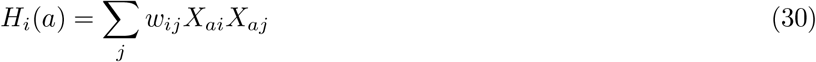

Statistical significance assessment is considered optional in this case, as it is possible to apply the above formula to just visualize the scores in space without filtering out non-significant values. However, an empirical approach has also been implemented for the purpose of avoiding considering interactions with low signal, and therefore visualizing only those cells or spots with significant scores. In this setting, if genes *a* and *b* are different, cell IDs in the counts matrix corresponding to gene *a* (or *b*) are shuffled *M* times (*M* = 1, 000 by default) and eventually, the most conservative p-values are selected. If gene *a* = *b*, though, the cell IDs in the counts matrix are shuffled. Then, the correlation values for each gene according to Eq. 29 and 30 are computed in each iteration, respectively, and Eq. 15 is used to calculate the p-value. These p-values are then adjusted using the Benjamini-Hochberg method [143].

Apart from using the empirical test, selecting the cells or spots based on the relative *H* scores is also a good option, where threshold values such as 1 standard deviation above the mean are especially recommended.

#### Test statistic 8: Is metabolite *m* exchanged by cell *i* and other spatially proximal cells?

Visualizing the individual cell/spot contributions of the results obtained using the metabolite-based approach described in **Test statistic 3** is also useful. For this, the equation below can be used for every metabolite (or ligand-receptor pathway), where the scores of the gene pairs (defined in Eq. 29 and 30) that are associated with a given metabolite are summed up:

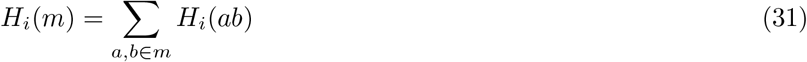

To test for statistical significance, an empirical test has been implemented. Here, Eq. 31 is computed in every shuffling iteration. This is done separately for genes *a* and *b* because, when *a* and *b* are different, the cell IDs corresponding to the counts matrices of both gene *a* and *b* are shuffled, giving rise to *H*_*i*_(*m*)(*a*) and *H*_*i*_(*m*)(*b*). However, if gene *a* = *b*, then the same output is considered for both *H*_*i*_(*m*)(*a*) and *H*_*i*_(*m*)(*b*). Then, two sets of p-values are obtained through Eq. 15, and eventually, the most conservative p-values are selected. P-values are also adjusted using the Benjamini-Hochberg procedure [143].

Apart from using the empirical test, selecting the cells or spots based on the relative *H* scores is also a good option, where threshold values such as 1 standard deviation above the mean are especially recommended.

#### Test statistic 9: Do genes *a* and *b* interact when *a* is expressed by cell *i* (that belongs to cell type *t*) and *b* by spatially nearby cells (that belong to cell type *u*)?

In addition to visualizing the cell-type-agnostic gene pair scores (defined in **Test statistic 7**), the equation below represents the individual cell/spot values when cell-type identity is considered:

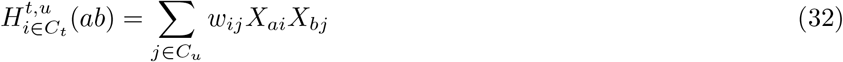

This equation is only used for visualization purposes, and no statistical test has been implemented.

#### Test statistic 10: Is metabolite *m* exchanged by cell *i* (that belongs to cell type *t*) and other spatially proximal cells (that belong to cell type *u*)?

In an analogous way to Eq. 32, the equation below represents the individual cell/spot metabolite (or ligand-receptor pathway) values when cell type identity is considered:

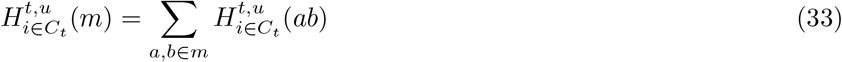

This equation is only used for visualization purposes, and no statistical test has been implemented.

### Null models for gene expression

To compute spatial correlation, we require a model for each gene and cell that defines the expected distribution of expression values under the null hypothesis of independent sampling. The null models described below are equivalent to the ones introduced in the Hotspot manuscript [35], which account for per-barcode library size and adjust the expected expression levels accordingly.

#### The negative binomial model

The negative binomial distribution is utilized to evaluate the significance of *H*. This distribution is commonly used in the single-cell RNA-seq field to model the counts that have been generated from experiments. It is defined as follows:

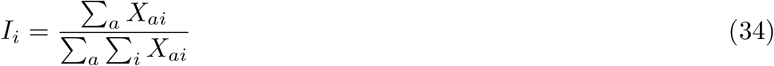

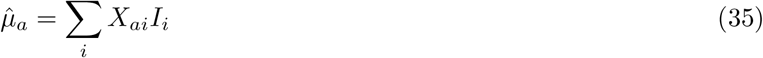

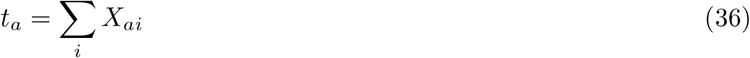

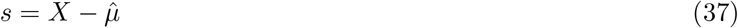

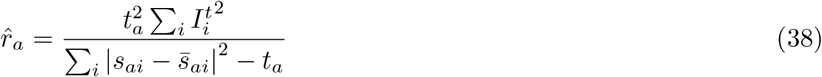

In this setting, *X*_*ai*_ and *t*_*a*_ represent the UMI count for gene *a* in cell *i* and the total number of UMI counts for gene *a*, respectively, and 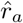 and *t*_*a*_ are the mean and dispersion of the negative binomial distribution. The expectation and variance of the expression values are estimated as:

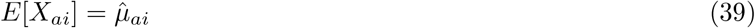

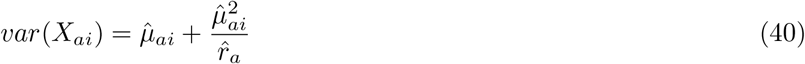

#### The Bernoulli model

Apart from the previously mentioned negative binomial model, the Bernoulli model can also be used, which estimates the detection of a gene (defined as a UMI count greater than 0). This model might be a better choice in case the data is very sparse. The model for gene *a* is as follows:

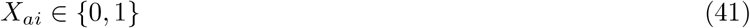

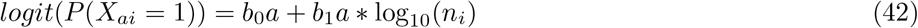

where *n*_*i*_ corresponds to the total number of UMI counts in cell *i*. Firstly, cells are aggregated into 30 bins based on the UMI distribution, where 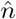 represents the bin center, and the average probability of detection per bin is computed as 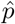 (cells with a UMI count of 0 are assigned a value of 1*e* − 10). Then linear regression is used to estimate model coefficients as 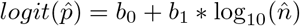.

### Feature elimination and selection

When assessing cellular crosstalk, it is useful to constrain the set of genes that will be used for inference to speed up computation time and reduce the number of hypotheses that will be generated. Before selecting relevant genes, those that are not expressed in more than a given fraction of the cells or spots (80% by default) can be eliminated. This is done to avoid performing feature selection on genes that are not relevant for cell-cell communication inference.

Then, different filtering strategies can be used to limit communication assessment to the most informative set of genes. The main method is local autocorrelation, which is used to select genes that spatially vary in a non-random manner, where only the genes that are significantly autocorrelated are selected for cell-cell communication assessment. For this, the approach developed in the Hotspot method [35] is used (**Test statistic 1**).

Apart from the local autocorrelation approach, for crosstalk inference using the cell type information, genes that (1) have a high enough expression and/or (2) are differentially overexpressed are selected for every cell type. For this, the following approach has been designed: on the one hand, for every cell type, genes that have a higher expression than 0 counts in at least 20% of the cells are selected. On the other hand, the Mann-Whitney U test is performed, and the genes that are significantly overexpressed (*FDR <* 0.05 and Cohen’s d *>* 0) with respect to the rest of the cells are kept. Finally, the genes that are selected in one or both filters are considered for each cell type.

### Computing gene pairs

Firstly, we select those genes that are associated with at least one metabolite or an LR pair and have passed the optional filters explained in the previous section. If cell type information is being used, gene pairs are computed for each cell type pair. For this, the given genes need to be relevant in their corresponding cell types. To compute gene pairs, the process slightly differs for LR pairs and metabolites. On the one hand, LR pairs are directly computed from CellChatDB (a pair is made up of a ligand and a receptor). On the other hand, for a given metabolite, all the possible pairwise combinations of the genes that carry that metabolite are considered. Only gene pairs in which both genes are confidently labeled as either importers or exporters are discarded. This is done to avoid inferring crosstalk between two importers or exporters and to try to maximize the cases where we find import-export interactions. For both metabolite and LR crosstalk, heterodimer information is taken into account so that all the components that make up a transporter (or a ligand or receptor) are jointly considered in the gene pair.

### Representing the equations as matrix multiplications

To efficiently compute the results, we represented the cell correlation as well as significance assessment formulas corresponding to the theoretical test as matrix multiplications. In all the equations, *X* is the gene expression matrix of dimension genes x cells and *W* is the weights matrix of dimension cells x cells. Δ_*t,u*_ is a matrix of dimension cells x cells with a value of 1 if a given cell pair belongs to cell types *t* and *u*, respectively, and 0 otherwise. *X*_1_ and *X*_2_ are the gene expression matrices of dimension ligands (or transporters) x cells and receptors (or transporters) x cells, respectively.

#### Test statistic (*H*) formulas

*Test statistic 1: Is gene a spatially autocorrelated? (Eq. 3)*

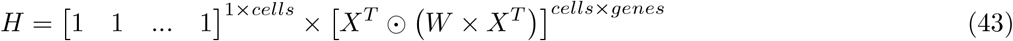

If *a* = *b* and gene *a* is involved in intercellular communication:

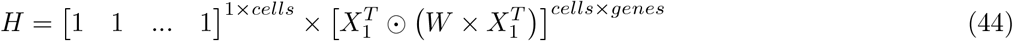

where the output *H* is a vector of length equal to the number of genes.

*Test statistic 2: Are genes a and b spatially co-localized (or interacting with each other)? (Eq. 16)*

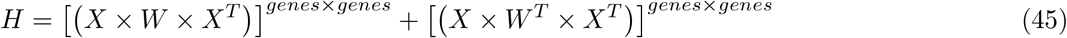

If gene pair (*a, b*) is involved in intercellular communication:

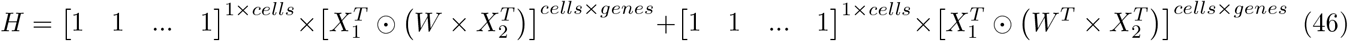

where the output *H* is a matrix of dimension genes x genes.

*Test statistic 5: Do genes a and b interact when expressed by cell types t and u, respectively? (Eq. 26)*

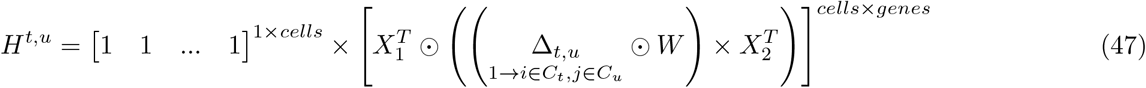

where the output *H*^*t,u*^ is a vector of length equal to the number of gene pairs.

*Test statistic 7: Do genes a and b interact when a is expressed by cell i and b by spatially nearby cells? (Eq. 29 and 30)*

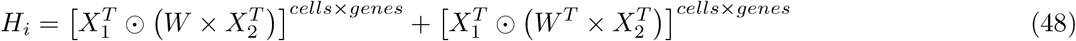

If a = b:

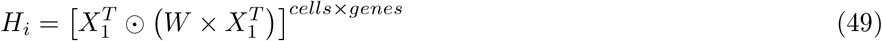

where the output *H*_*i*_ is a matrix of dimension cells x genes.

*Test statistic 9: Do genes a and b interact when a is expressed by cell i (that belongs to cell type t) and b by nearby cells (that belong to cell type u)? (Eq. 32)*

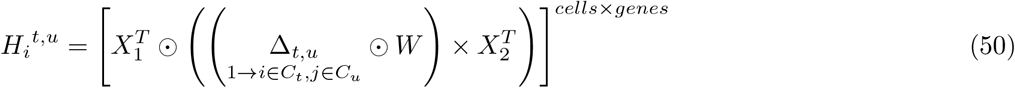

where the output *H*_*i*_^*t,u*^ is a matrix of dimension cells x genes.

#### Formulas corresponding to the second moments of *H*

*Test statistic 2: Are genes a and b spatially co-localized (or interacting with each other)? (Eq. 24)*

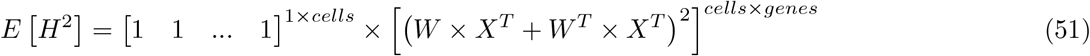

If gene pair (*a, b*) is involved in intercellular communication:

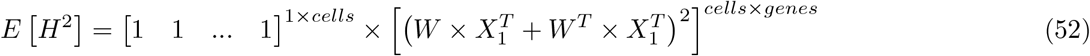

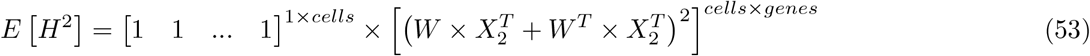

where the output *E*[*H*^2^] is a vector of length equal to the number of genes.

### Data analysis

#### Running Harreman on the human lung data

To test Harreman on high-resolution spatial data, we applied it on a human renal cell carcinoma, post anti-PD1, lung metastasis dataset made available by *Liu et. al*. [44]. In this study, the authors introduced Slide-TCR-seq, where they sequenced joint transcriptomic and T cell receptor (TCR) information across diverse tissues and samples. Raw counts were normalized using the *sc*.*pp*.*normalize_total* function with a target sum of 10,000, and we then applied the *sc*.*pp*.*log1p* function to log-transform the normalized counts. This was done separately for each slide. All the slides were collected from a single patient and were sequenced in two different batches.

Firstly, Harreman was run on all the 7 samples together (Harreman both computes the proximity graph and scales the gene expression values separately for each sample) to infer the metabolic zonation of the tissue (Fig. 2B and Fig. S3). For this, the PyTorch version of the Hotspot algorithm [35] was used by only considering metabolic genes obtained from the Compass algorithm [130] (**Test statistics 1 and 2**). Prior to running the analysis, those genes expressed in less than 50 spots in all the slides were filtered out. The functions and their corresponding parameters used for this analysis are as follows: the *harreman*.*tl*.*compute_knn_graph* function with *compute_neighbors_on_key=“spatial”, neighborhood_radius=100, weighted_graph=False*, and *sample_key=‘sample’* ; the *harreman*.*hs*.*compute_local_autocorrelation* function with *layer_key=“counts”, model=‘bernoulli’, species=‘human’*, and *use_metabolic_genes=True*. Pairwise correlations were computed for all the significantly autocorrelated genes (theoretical FDR < 0.01) using the *harreman*.*hs*.*compute_local_correlation* function. To create gene modules, the *min_gene_threshold=20* parameter was used in the *harreman*.*hs*.*create_modules* function, ending up with a total of 4 modules shared across all slides. To compare the theoretical and empirical test results, the *permutation_test=True* parameter was used in the *harreman*.*hs*.*compute_local_autocorrelation* and *harreman*.*hs*.*compute_local_correlation* functions (Fig. S2A; first and second figures starting from the left).

To assign spots as top-scoring for a module (Fig. 2C), the *harreman*.*hs*.*compute_top_scoring_modules* function was used with the *sd=1* and *use_super_modules=False* parameters, where we z-normalized all module scores across spots, and then required a top-scoring spot to very specifically express a given module and not express any others. To ensure this, we required top-scoring spots to have (1) a score of >1 standard deviation (s.d.) above its mean module score, and (2) a score of <1 s.d. above the mean for all other modules. If the first condition is not met, that is, if none of the modules has a value higher than 1 s.d. above the mean for a given spot, then the module with the highest z-normalized value is assigned to the spot. In case the second condition is not met, that is, if there is more than one module with a value higher than 1 s.d. above the mean for a given spot, then the module with the highest z-normalized value among those modules that meet condition (1) is selected. Finally, if none of the modules have a higher value than the mean for a given spot, then this spot ends up unassigned.

Then, to evaluate the metabolite exchange as well as the ligand-receptor interactions in the tissue (**Test statistics 3 and 2**, respectively; Fig. 3), the same neighborhood graph as in the metabolic zonation analysis (**Test statistics 1 and 2**) was used, but before performing the interaction analysis, the *harreman*.*tl*.*apply_gene_filtering* function with the *model = ‘bernoulli’* and *autocorrelation_filt = True* parameters was used to keep genes that were spatially autocorrelated (theoretical FDR < 0.05; **Test statistic 1**). Gene pairs were computed using the *harreman*.*tl*.*compute_gene_pairs* function (*ct_specific = False*). The cell-cell communication analysis (*harreman*.*tl*.*compute_cell_communication* function) was performed using the *bernoulli* model (*model = ‘bernoulli’*) to standardize the raw counts, and the log-normalized counts were used for the empirical test (*layer_key_np_test = ‘log_norm’*). Further, both the theoretical and empirical tests were performed (*test = “both”*), using 1000 permutations for the empirical test (*M = 1000*), allowing us to compare the theoretical and empirical test results (Fig. S2A; third and fourth figures starting from the left). Eventually, significant communication events (empirical FDR < 0.05) were considered (**Test statistics 2 and 3**). For this, the *harreman*.*tl*.*select_significant_interactions* function with the *test = “non-parametric”* and *threshold = 0*.*05* parameters was used. We then computed metabolite and ligand-receptor pathway scores for each spot using the log-normalized gene expression values (**Test statistic 8**; Fig. S5). This was done using the *harreman*.*tl*.*compute_interacting_cell_scores* function with the *test = “both”* parameter. Only those gene pairs and metabolites that were labeled as statistically significant in the previous analysis were considered.

To group metabolites with a similar spatial profile together, we binarized the **Test statistic 8** scores by assigning a value of 1 if the value of a given spot is higher than 1 standard deviation above the mean and 0 otherwise. Eventually, taking these binarized scores, we ran the pairwise correlation analysis using the *‘bernoulli’* model (**Test statistic 4**). The same parameters as before were used to create the neighborhood graph, and those metabolites with all-zero values in at least one of the samples were filtered out for subsequent pairwise correlation. Eventually, modules were created, and after grouping similar ones, we ended up with 4 interpretable metabolite groups (Fig. 3A).

Finally, some of the most interesting metabolites and their respective significant gene pairs were selected to perform a cell type-aware analysis (**Test statistics 5 and 6**; Fig. 3F). For this, the cell type with the highest z-normalized DestVI-inferred proportion was assigned to each spot (deconvolved counts were also used for cell type-aware inference, but the results weren’t considered for downstream analysis because the results that we got were too general; that is, we got significant results for most of the cell type pairs). First, gene pairs were computed for each cell type pair using the *harreman*.*tl*.*compute_gene_pairs* function with the *cell_type_key=‘cell_type’* parameter. Then, the cell type-aware crosstalk analysis was performed using the *harreman*.*tl*.*compute_ct_cell_communication* function with the following parameters: *model=‘bernoulli’, cell_type_key=‘cell_type’, M = 1000, test = “both”, layer_key_p_test=‘counts’, layer_key_np_test=‘log_norm’, subset_gene_pairs=gene_pairs, subset_metabolites=metabolites*, and *fix_gp=fix_gp*, where *gene_pairs* and *metabolites* represent the gene pairs and metabolites of interest, respectively. This analysis was performed in two different ways. On the one hand, we performed communication inference by shuffling gene expression of cells within the cell type pair of interest (cell type identity and spatial weights are kept constant; *fix_gp=False*). Through this approach, we aim for gene pairs (or metabolites) that are co-expressed (or where the observed co-expression of all gene pairs associated with a given metabolite) by their corresponding cell types more than expected by chance (**Test statistic 5 (or 6)**; empirical null hypothesis 1). The second approach helps us answer the following question: Which cell types are driving a given gene pair’s spatial pattern (or the observed co-expression of all gene pairs associated with a given metabolite)? Therefore, what we do in this case is shuffle the cell type identity by keeping the spatial weights and gene expression counts constant (**Test statistic 5 (or 6)**; empirical null hypothesis 2; *fix_gp=True*). We then focused on the cell type pairs labeled as significant (FDR < 0.01) in the first hypothesis. The parameters needed to create the spatial proximity graph are the same ones used for the previous analyses. To visualize the cell type-specific scores for every spot, the *harreman*.*tl*.*compute_ct_interacting_cell_scores* function was used.

#### Running Harreman on the mouse colon data

To study the metabolic differences across conditions, the mouse colon 10x Visium dataset curated by *Parigi et. al*. was used [96]. In this study, the authors characterized the transcriptomic profile of the steady state and healing mouse colon following inflammation induced by Dextran Sulfate Sodium (DSS). The digital unrolling corresponding to the day 14 sample was performed by adapting the code written for the day 0 condition (https://github.com/ludvigla/healing_intestine_analysis/blob/main/doc/Digital_unrolling.Rmd).

To determine the metabolic zonation of the tissue, Harreman was run on the unrolled space of both conditions at the same time by computing the proximity graph and scaling the gene expression values separately for each condition (Fig. 4C and Fig. S7). For this, the optimized version of the Hotspot algorithm [35] was used by only considering metabolic genes obtained from the Compass algorithm [130] (**Test statistics 1 and 2**). Before running the analysis, those genes expressed in fewer than 50 spots in both conditions were filtered out. The functions and their corresponding parameters used for this analysis are as follows: the *harreman*.*tl*.*compute_knn_graph* function with *compute_neighbors_on_key=“spatial_unrolled”, n_neighbors=5, weighted_graph=False*, and *sample_key=‘cond’* ; the *harreman*.*hs*.*compute_local_autocorrelation* function with *layer_key=“counts”, model=‘danb’, species=‘mouse’*, and *use_metabolic_genes=True*. Pairwise correlations were computed for all the significantly autocorrelated genes (theoretical FDR < 0.01) using the *harreman*.*hs*.*compute_local_correlation* function. To create gene modules, the *min_gene_threshold=20* parameter (default value) was used in the *harreman*.*hs*.*create_modules* function. 10 modules were created (Fig. S7A,B), but five of them were eventually merged into two super-modules, ending up with a total number of 7 super-modules present across both slides. This decision was made after (1) getting a highly positive Pearson correlation of the module scores and (2) ensuring the local correlations between genes of grouped modules were high on average. The super-modules were labeled, and those with the same label were merged. Module scores were computed both for the final super-modules by using the *harreman*.*hs*.*calculate_super_module_scores* function. To compare the theoretical and empirical test results, the *permutation_test=True* parameter was used in the *harreman*.*hs*.*compute_local_autocorrelation* and *harreman*.*hs*.*compute_local_correlation* functions (Fig. S2B; first and second figures starting from the left).

To compare the obtained modules with the ones created by running the metabolic zonation pipeline on the original spatial coordinates (Fig. S8), the same functions and parameters as in the previous paragraph were used except the following: *compute_neighbors_on_key=“spatial”* (instead of *compute_neighbors_on_key=“spatial_unrolled”*) and *neighborhood_radius=150* (instead of *n_neighbors=5*).

To assign spots as top-scoring for a super-module (module from now on; Fig. 4D), the *harreman*.*hs*.*compute_top_scoring_modules* function was used with the *sd=1* and *use_super_modules=True* parameters, where we z-normalized all module scores across spots, and then required a top-scoring spot to very specifically express a given module and not express any others. To ensure this, the same procedure as in the previous subsection for the human lung dataset was used.

Despite all modules being assigned to at least one spot in both conditions, we wanted to evaluate if there are significant differences in the module scores between both conditions for every module. For this, we performed a Mann-Whitney U test and eventually adjusted the p-values using the Benjamini-Hochberg approach [143].

To dig deeper into the tissue localization of the modules along the proximal-distal and serosa-luminal axes, the coordinates from the unrolled space were used. For this, the z-scored Hotspot module scores were smoothed using a polynomial regression model with bootstrap resampling (1000 iterations).

Then, to evaluate the metabolite exchange as well as the ligand-receptor interactions in the tissue (**Test statistics 3 and 2**, respectively; Fig. 5), the same neighborhood graph as in the metabolic zonation analysis (**Test statistics 1 and 2**) was used, but prior to performing the interaction analysis, the *harreman*.*tl*.*apply_gene_filtering* function with the *model = ‘danb’* and *autocorrelation_filt = True* parameters was used to keep genes that were spatially autocorrelated (theoretical FDR < 0.05; **Test statistic 1**). Gene pairs were computed using the *harreman*.*tl*.*compute_gene_pairs* function (*ct_specific = False*). The cell-cell communication analysis (*harreman*.*tl*.*compute_cell_communication* function) was performed using the *danb* model (*model = ‘danb’*) to standardize the raw counts, and the log-normalized counts were used for the empirical test (*layer_key_np_test = ‘log_norm’*). Further, both the theoretical and empirical tests were performed (*test = “both”*), using 1000 permutations for the empirical test (*M = 1000*), allowing us to compare the theoretical and empirical test results (Fig. S2B; third and fourth figures starting from the left). Eventually, significant communication events (empirical FDR < 0.05) were considered (**Test statistics 2 and 3**). For this, the *harreman*.*tl*.*select_significant_interactions* function with the *test = “nonparametric”* and *threshold = 0*.*05* parameters was used. We then computed metabolite and ligand-receptor pathway scores for each individual spot using the log-normalized gene expression values (**Test statistic 8**; Fig. S10). This was done using the *harreman*.*tl*.*compute_interacting_cell_scores* function with the *test = “both”* parameter. Only those gene pairs and metabolites that were labeled as statistically significant in the previous analysis were considered.

To group metabolites according to their spatial pattern, we binarized the **Test statistic 8** scores by assigning a value of 1 if the value of a given spot is higher than 1 standard deviation above the mean and 0 otherwise. Eventually, taking these binarized scores, we ran Hotspot using the *‘bernoulli’* model (**Test statistic 4**). The same parameters as before were used to create the neighborhood graph, and those metabolites with all-zero values in at least one of the samples were filtered out for subsequent pairwise correlation. 5 metabolite groups were computed after grouping spatially similar ones (Fig. 5A).

To perform the finest analysis of the Harreman pipeline, that is, the cell-type-aware analysis, some of the most interesting metabolites and their respective significant gene pairs were selected (**Test statistics 5 and 6**; Fig. 3G). For this, the procedure explained in the previous subsection for the human lung dataset was implemented. For this, the cell type with the highest z-normalized DestVI-inferred proportion was assigned to each spot (deconvolved counts were also used for cell type-aware inference, but the results weren’t considered for downstream analysis because the results that we got were too general; that is, we got significant results for most of the cell type pairs). First, gene pairs were computed for each cell type pair using the *harreman*.*tl*.*compute_gene_pairs* function with the *cell_type_key=‘cell_type’* parameter. Then, the cell type-aware crosstalk analysis was performed using the *harreman*.*tl*.*compute_ct_cell_communication* function with the following parameters: *model=‘danb’, cell_type_key=‘cell_type’, M = 1000, test = “both”, layer_key_p_test=‘counts’, layer_key_np_test=‘log_norm’, subset_gene_pairs=gene_pairs, subset_metabolites=metabolites*, and *fix_gp=fix_gp*, where *gene_pairs* and *metabolites* represent the gene pairs and metabolites of interest, respectively. This analysis was performed in two different ways, as explained in the previous subsection for the human lung dataset. The parameters needed to create the spatial proximity graph are the same ones used for the previous analyses. To visualize the cell type-specific scores for every spot, the *harreman*.*tl*.*compute_ct_interacting_cell_scores* function was used.

Finally, some of the findings obtained in the murine tissue were translated into a human single-cell dataset [109] by performing a differential expression analysis (using the *sc*.*tl*.*rank_genes_groups* function [145] with the *method=“wilcoxon”* parameter) between spots exchanging either vitamin A or LPC and those not exchanging any of them. For this, we focused on the distal part of the day 14 tissue (the region delineated by the immune modules, that is, Modules 5-7). Then, we built a gene signature by selecting the top 200 differentially expressed genes (LFC > 0 and FDR < 0.05; sorted by LFC) and computed the signature scores (as explained in the “Signature score computation” subsection below) in the human dataset. For each disease group (ulcerative colitis (UC) and Crohn’s disease (CD)) and cellular compartment (epithelium, immune, and stroma), the bottom and top 20% of the signature score distribution were labeled as “Low” and “High”, respectively. We then visualized the inflammation score difference between these two groups for each case, and a Mann-Whitney U test was performed to assess statistical significance (Fig. 5J).

#### Robustness of the metabolic zonation analysis

We further assessed the robustness of the metabolic zonation analysis to the varying neighborhood radius (human lung dataset) or the number of neighbors (mouse colon dataset) used to create the spatial proximity graph to analyze its effect on the gene selection based on autocorrelation and on module creation. For the first case, we performed the gene autocorrelation analysis (**Test statistic 1**) keeping the same parameters but using a different neighborhood radius (human lung dataset) or number of nearest neighbors (mouse colon dataset) as input (with a radius of 50–400 *µ*m in increments of 50 *µ*m; or 5-50 nearest neighbors in increments of 5 neighbors). We then calculated for each gene the difference in autocorrelation score against its autocorrelation when using 100 *µ*m (the reference value; or 5 neighbors), and we saw that this difference is minimal across the diverse runs using different values for the spatial proximity graph (Fig. S4A and Fig. S9A).

To evaluate the module similarity across neighborhood radius (or number of neighbors) values, we computed the Jaccard similarity between each original module and the most similar modules generated using a different value of neighborhood radius (or number of neighbors). We visualized this distribution as a box plot (Fig. S4B and Fig. S9B). We also computed the Pearson correlation between the inferred modules in the dataset and the most similar modules (based on maximum correlation coefficient) generated using a different number of of neighborhood radius (or number of neighbors). We also visualized this distribution as a box plot (Fig. S4E and Fig. S9E). We also applied the same procedure across different minimum number of gene thresholds for module creation (with a value of 10–50 genes in increments of 5 genes, with the reference value being 20 genes; Fig. S4C,F and Fig. S9C,F). This experiment shows that, in general, the modules created using the reference values are robust to the varying number of neighbors as well as the minimum gene threshold.

Finally, we wanted to evaluate the relationship between pairwise and Pearson correlation. This analysis showed a high correlation between the Pearson R values and the pairwise correlation (**Test statistic 2**) Z-scores (Fig. S4D and Fig. S9D). Further, despite the global correlation approach being able to also find a similar module structure (Fig. S4G and Fig. S9G), the signal was weaker compared to the pairwise correlation Z-scores obtained from our approach (Fig. S4H and Fig. S9H), confirming the sensitivity of pairwise correlation by constraining the analysis to local neighborhoods.

#### Deconvolution of the spatial data

To assign cell type information to each spot, DestVI [47] was used. For this, in the human lung dataset, the same reference single-cell dataset [46] used in the original paper was used, while for the mouse colon dataset, a single-cell dataset from an unpublished project was used. From the single-cell dataset, genes with a total number of UMI counts less than 10 were filtered out, and the 8000 most highly variable genes, as well as the transporters, ligands, and receptors that are part of both HarremanDB and CellChatDB, were then used to train the scLVM. Genes that are present both in the single-cell and spatial datasets were kept for deconvolution (6,047 and 8,320 in the lung and colon datasets, respectively). After this model was trained, all spatial slides were deconvoluted together. Therefore, DestVI was run once to obtain the deconvolution results of all slides.

The parameters that we used in the *CondSCVI* function were *weight_obs=False, prior=‘mog’*, and *num_classes_mog=10*. The sample ID variable was used as batch key in the *CondSCVI*.*setup_anndata* function. To train the single cell model, a batch size of 1024 and a maximum epoch number of 2000 were used. To perform deconvolution using the spatial latent variable model, the parameters that were used in the *DestVI*.*from_rna_model* function were *add_celltypes=2, n_latent_amortization=None, anndata_setup_kwargs={‘smoothed_layer’: ‘smoothed’}*, and *prior_mode = ‘normal’*. To train the spatial model, the parameters used were as follows: *max_epochs=1000, n_epochs_kl_warmup=200, batch_size=1024*, and *plan_kwargs={‘weighting_kl_latent’: 1e-2, ‘ct_sparsity_weight’: 0}*. The ‘smoothed’ layer of the AnnData was created by computing the matrix multiplication between the raw counts and the spatial neighborhood connectivity matrix. The latter was computed by using the *sc*.*pp*.*neighbors* function with 5 neighbors.

To estimate the relative abundance of cell types across samples, the cell type proportion information provided by DestVI was z-scored, and the average value was computed for each module and sample (Fig. 2E and Fig. 4F).

#### Cell type-specific differential expression analysis

To study the differences in effector phenotype of different cell types across metabolic modules, a module-vs-all DE analysis was performed, assessing the differential expression of a given cell type in one module compared to the rest of the modules. The cell types considered for the analysis were macrophages and T-Helper cells (CD4+ T cells) from the human lung dataset [44] (Fig. 2F-H). For this, the *de_genes* function of the *destvi_utils* package was used (https://github.com/YosefLab/destvi_utils) [47], which makes use of the DestVI-generated cell type-specific gene expression data. For each cell type, the spots with a DestVI-inferred cell type proportion higher than 1 standard deviation above the mean were considered for the analysis.

To identify differences in the expression profile of cells that belong to a given cell type and exchange a given metabolite of interest versus the ones that don’t, the same procedure as above was used. However, for each cell type, the spots that were labeled with the cell type of interest were considered for the analysis (Fig. 3H and Fig. 5I).

#### Differential metabolite score analysis

To find condition-specific metabolites in the mouse colon dataset [96], the Mann-Whitney U test between the day 14 and day 0 conditions was performed on the min-max normalized metabolite scores from the non-parametric test (Fig. 5D; **Test statistic 8**). Only the scores corresponding to the spots with a value higher than 1 standard deviation above the mean were considered for the analysis (the scores corresponding to the rest of the spots were converted to 0). The Cohen’s d value was used to report the effect size.

#### Survival analysis

Given the spatial pattern of the *SLC8A1* transporter gene in the tumor boundary of the human lung dataset [44], survival analysis was performed using the GEPIA2 software [146]. For this, we focused on the Kidney renal clear cell carcinoma (KIRC) TCGA dataset, and patients were stratified into *SLC8A1* high and low groups (Fig. 3E). The same analysis was also performed on two lung tumors (Fig. S6A): Lung adenocarcinoma (LUAD) and Lung squamous cell carcinoma (LUSC).

#### Signature score computation

Signature scores of the lung and kidney were computed using the Python version of the VISION software [147] (visionpy; https://github.com/YosefLab/visionpy). visionpy calculates the signature scores as follows:

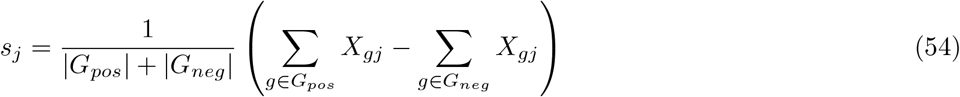

where *s*_*j*_ is the signature score in cell *j*; *X*_*gj*_ is the log-normalized expression of gene *g* in cell *j*; and *G*_*pos*_ and *G*_*neg*_ are the set of ‘positive’ and ‘negative’ signature genes, respectively. Due to the observation that, depending on the scaling or normalization method used, signature scores can still be highly correlated with cell-level metrics such as the number of UMIs per cell, signature scores are then Z-normalized by using the expected value and variance of a randomly-drawn signature in cell *j*.

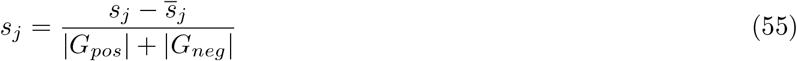

The expected value is defined as follows:

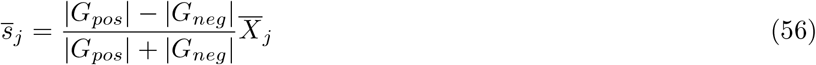

While the expected variance is:

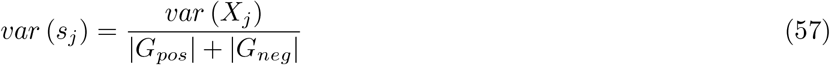

The signature scores for the following signatures were computed (hand-curated list; Fig. S6B):

1. Kidney: *CA9, VIM, PAX8, NDUFA4L2, EGLN3, ENO2, SLC16A3, SLC2A1, SLC8A1, AQP1, CP, FABP7, GGT1, SLC17A3, PLIN2, GPX3, NNMT*, and *PCSK1N*.
2. Lung: *NKX2-1, SFTPB, SFTPC, NAPSA, CLDN18, KRT5, TP63, SOX2, FGFR1, MUC1, TTF1, CEACAM5, EPCAM, KRT7, SCGB1A1, CDH1, FOXA2, GATA6*, and *B3GNT7*.

## Data availability

All datasets used in this study are publicly available except the single-cell reference used for deconvolution of the mouse colon dataset, which belongs to an unpublished work and is not publicly available yet. Raw counts and spatial coordinates of the mouse colon dataset were obtained from https://github.com/ludvigla/healing_intestine_analysis, GEO: GSE169749; the AnnData file corresponding to the human intestinal dataset (*TAURUS_raw_counts_annotated_final*.*h5ad*) was downloaded from Zenodo https://doi.org/10.5281/zenodo. 13768607; raw counts and spatial coordinates of the human lung dataset were obtained from https://cellxgene.cziscience.com/collections/02b01703-bf1b-48de-b99a-23bef8cccc81; the single-cell reference for deconvolution of the human lung dataset was obtained from https://singlecell.broadinstitute.org/single_cell/study/SCP1288/tumor-and-immune-reprogramming-during-immunotherapy-in-advanced-renal-cell-carcinoma#/.

For digital unrolling of the mouse colon data, tif images corresponding to both conditions were downloaded from GSM5213483 (day 0) and GSM5213484 (day 14) and were added to the *spatial* folder (under *healing_intestine_analysis/data/V19S23-097_A1* and *healing_intestine_analysis/data/V19S23-097_B1* for day 0 and day 14, respectively). The Spatial Slide-seq as well as the Single-cell PBMC CD4 datasets used to compare the original Hotspot algorithm with the PyTorch version of Harrreman were collected from https://github.com/YosefLab/scVI-data/blob/master/rodriques_slideseq.h5ad?raw=true and https://support.10xgenomics.com/single-cell-gene-expression/datasets/3.1.0/5k_pbmc_protein_v3_nextgem, respectively. For the latter dataset, the *5k_pbmc_protein_v3_filtered_feature_bc_matrix*.*h5* file (“Feature / cell matrix HDF5 (filtered)”) was used. Both datasets were already used in the Hotspot manuscript [35].

## Code availability

Harreman is available as an open source Python package at https://github.com/YosefLab/Harreman. All code to reproduce the analyses and the figures included in the manuscript is available at https://github.com/YosefLab/Harreman-reproducibility. We provide comprehensive documentation as well as tutorials at https://harreman.readthedocs.io/.

## Acknowledgments

We thank Can Ergen, Nathan Levy, Jonas Maaskola, Florian Ingelfinger, Artemii Bakulin, and Eli Benuck for valuable discussions and feedback on this project.

## Supplementary Figures

**Figure S1:**
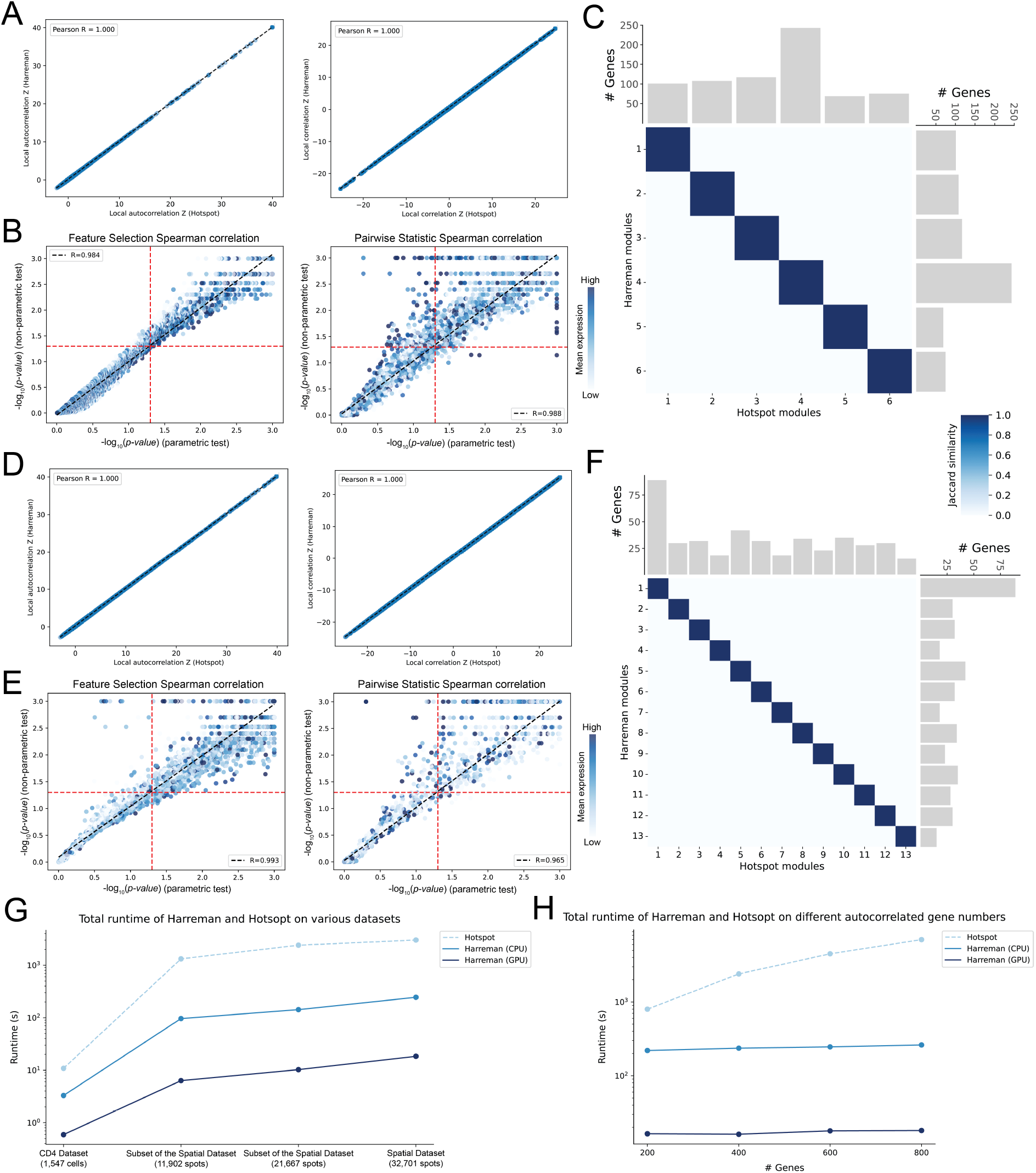
Comparison between the original and the optimized version of the Hotspot algorithm. *Spatial Slide-seq dataset (Puck_180819_12 sample)*: (A) Pearson correlation of the gene autocorrelation (**Test statistic 1**) as well as the pairwise correlation (**Test statistic 2**) Z-scores between the PyTorch Hotspot version implemented in Harreman and the original Hotspot algorithm. (B) Spearman correlation between the autocorrelation (left) or pairwise correlation (right) parametric and non-parametric test significance results. Mean expression represents the average log-normalized gene expression across spots that express a given gene. The dashed red lines represent the p-value = 0.05 threshold. (C) Jaccard similarity score between modules identified by Hotspot and modules identified by Harreman. *Single-cell PBMC CD4 dataset* : (D), (E), and (F) are equivalent to (A), (B), and (C), respectively. (G) Runtime comparison between Hotspot and Harreman (CPU and GPU) on the PBMC dataset, as well as different subsets of the Slide-seq dataset. (H) Runtime comparison between Hotspot and Harreman on the Slide-seq dataset using different numbers of significantly autocorrelated genes.

**Figure S2:**
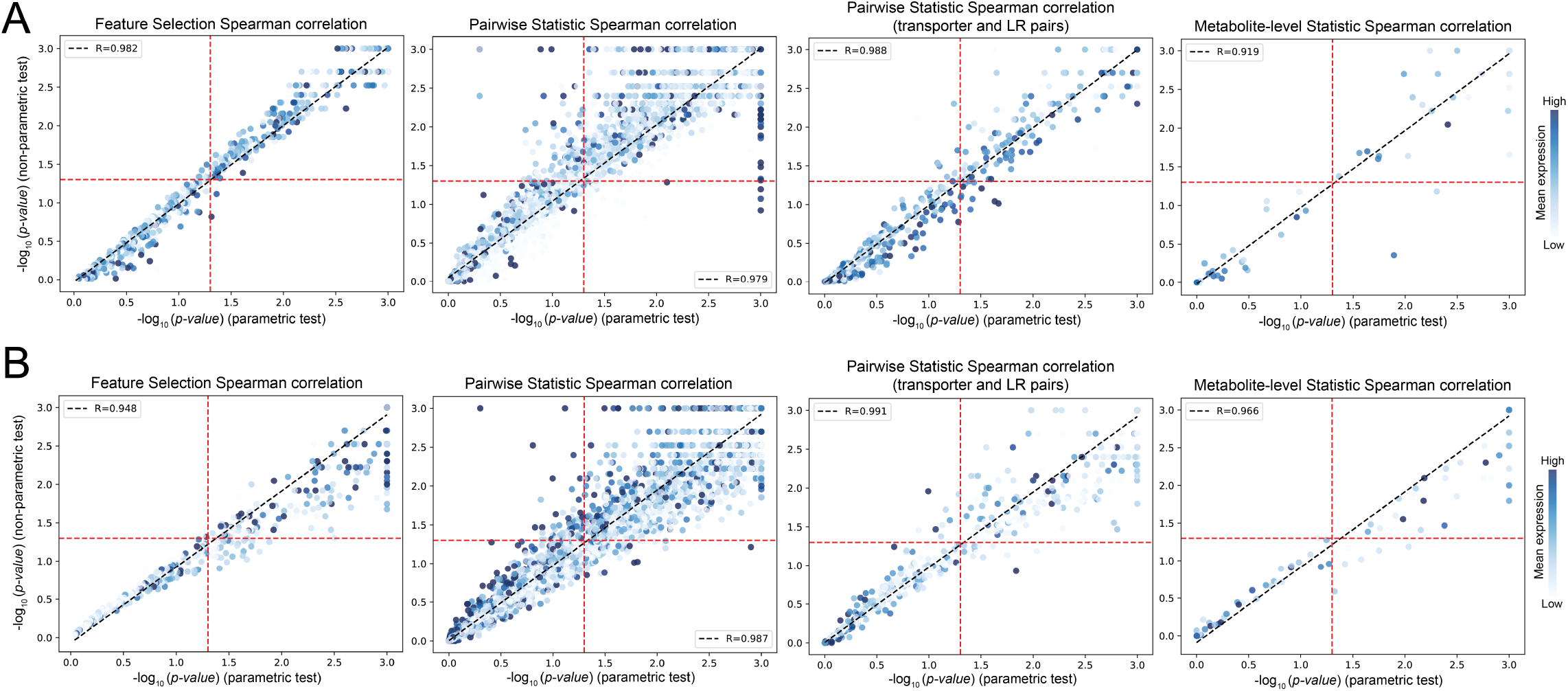
Comparison between the parametric and non-parametric test statistics. *Slide-seq human lung dataset:* (A) Correlation between the parametric and non-parametric test significance results for the following statistics: gene autocorrelation (all metabolic genes; **Test statistic 1**), pairwise correlation (metabolic genes selected from the autocorrelation test; **Test statistic 2**), pairwise correlation (transporters and ligand-receptor (LR) pairs selected from the autocorrelation test; **Test statistic 2**), and metabolite-level correlation (**Test statistic 3**). The dashed red lines represent the p-value = 0.05 threshold. *10x Visium mouse colon dataset:* (B) Equivalent to (A).

**Figure S3:**
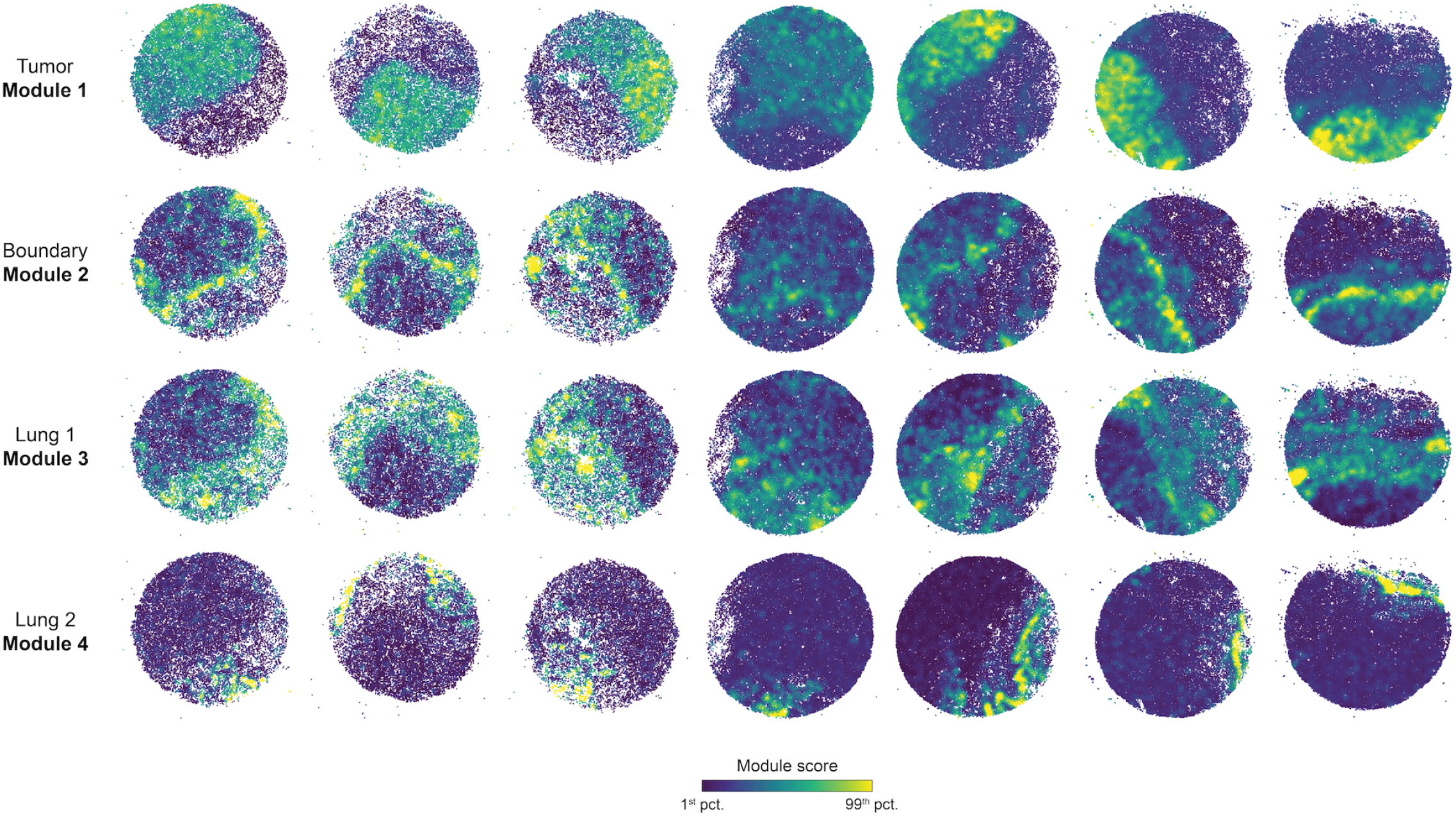
Metabolic zonation on the human lung dataset. Spatial plots of the metabolic gene module scores across samples.

**Figure S4:**
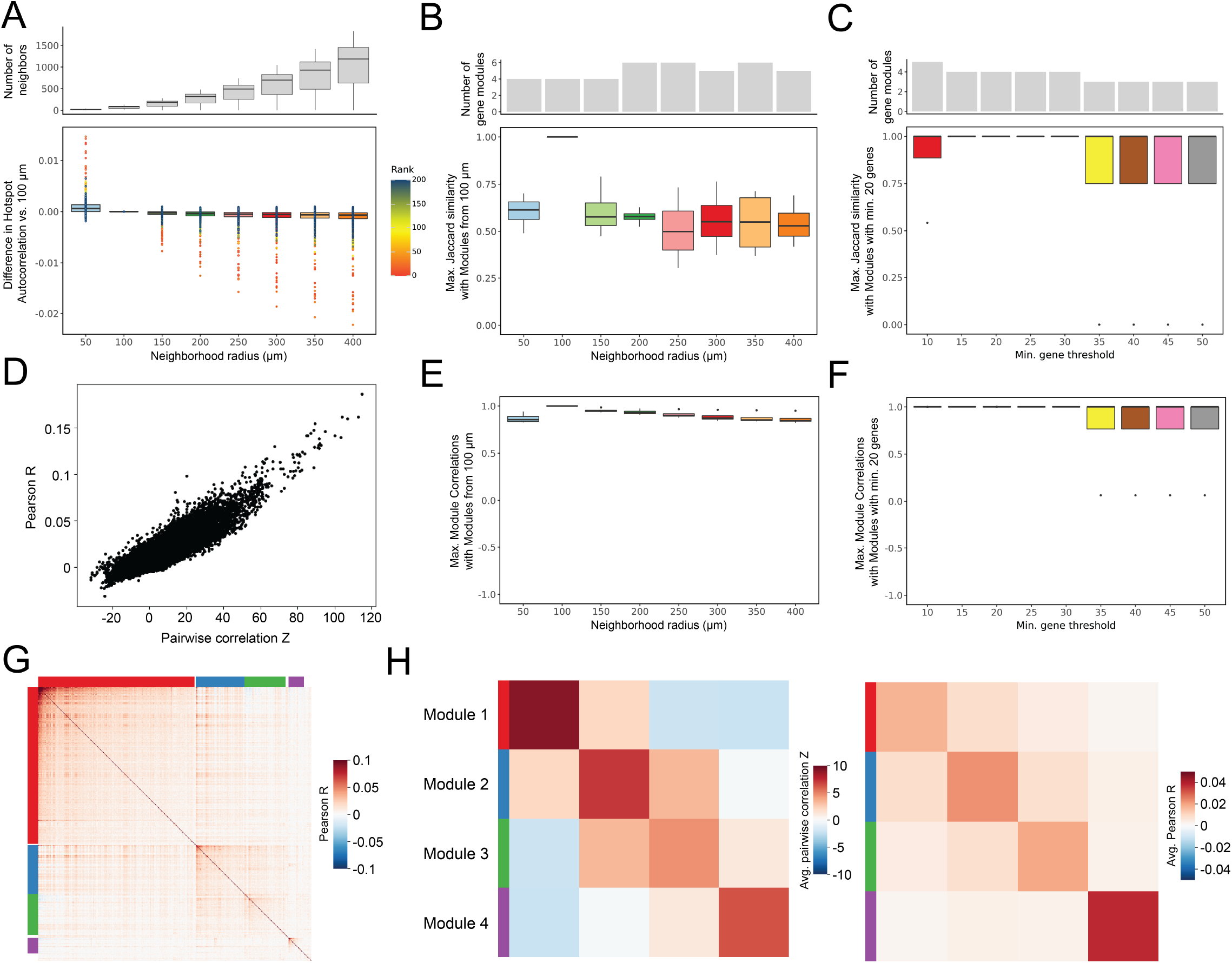
Gene autocorrelation and detected modules in the human lung dataset are robust to varying parameter values. (A) Top: Distribution of the number of neighbors across different neighborhood radius values. Bottom: Difference in gene autocorrelation (**Test statistic 1**) score between a neighborhood radius of 100 *µ*m and varying neighborhood radius values. (B) Top: Number of gene modules across varying neighborhood radius values. Bottom: Jaccard similarity between the inferred 4 modules in this dataset (neighborhood radius of 100 *µ*m) and the most similar modules generated using a different neighborhood radius value. (C) Top: Number of gene modules across varying minimum gene threshold values. Bottom: Jaccard similarity between the inferred 4 modules in this dataset (minimum gene threshold of 20 genes) and the most similar modules generated using a different minimum gene threshold value. (D) Comparison of Pearson correlation R and pairwise correlation (**Test statistic 2**) Z values for all significantly autocorrelated (theoretical FDR < 0.01; **Test statistic 1**) genes. (E) Pearson correlation between the inferred 4 modules in this dataset (neighborhood radius of 100 *µ*m) and the most similar modules generated using a different neighborhood radius value. (F) Pearson correlation between the inferred 4 modules in this dataset (minimum gene threshold of 20 genes) and the most similar modules generated using a different minimum gene threshold value. (G) Gene-gene Pearson correlation for the 605 genes selected using **Test statistic 1**. (H) Average pairwise correlation Z (left; **Test statistic 2**) and Pearson R (right) values.

**Figure S5:**
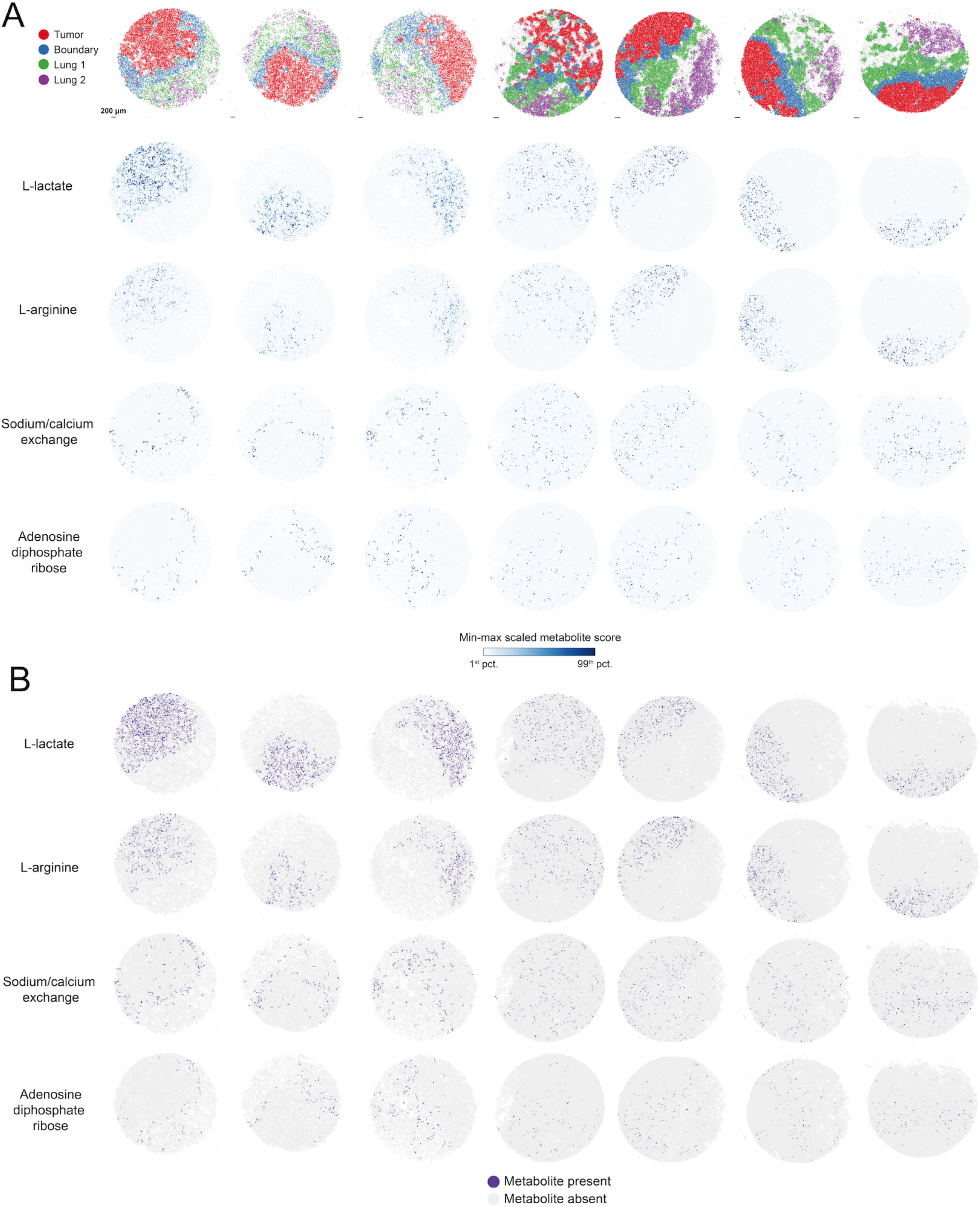
Metabolite abundance across samples in the human lung dataset. (A) Top: Spatial distribution of the modules across samples. Scale bars, 200 *µ*m. Bottom: Spatial plots of the min-max scaled metabolite scores (**Test statistic 8**) for some of the region-specific metabolites. (B) Spatial plots of the binarized metabolite scores used for metabolite module grouping (**Test statistic 4**). Those scores with a Z-scored value higher than 1 standard deviation above the mean were labeled as present.

**Figure S6:**
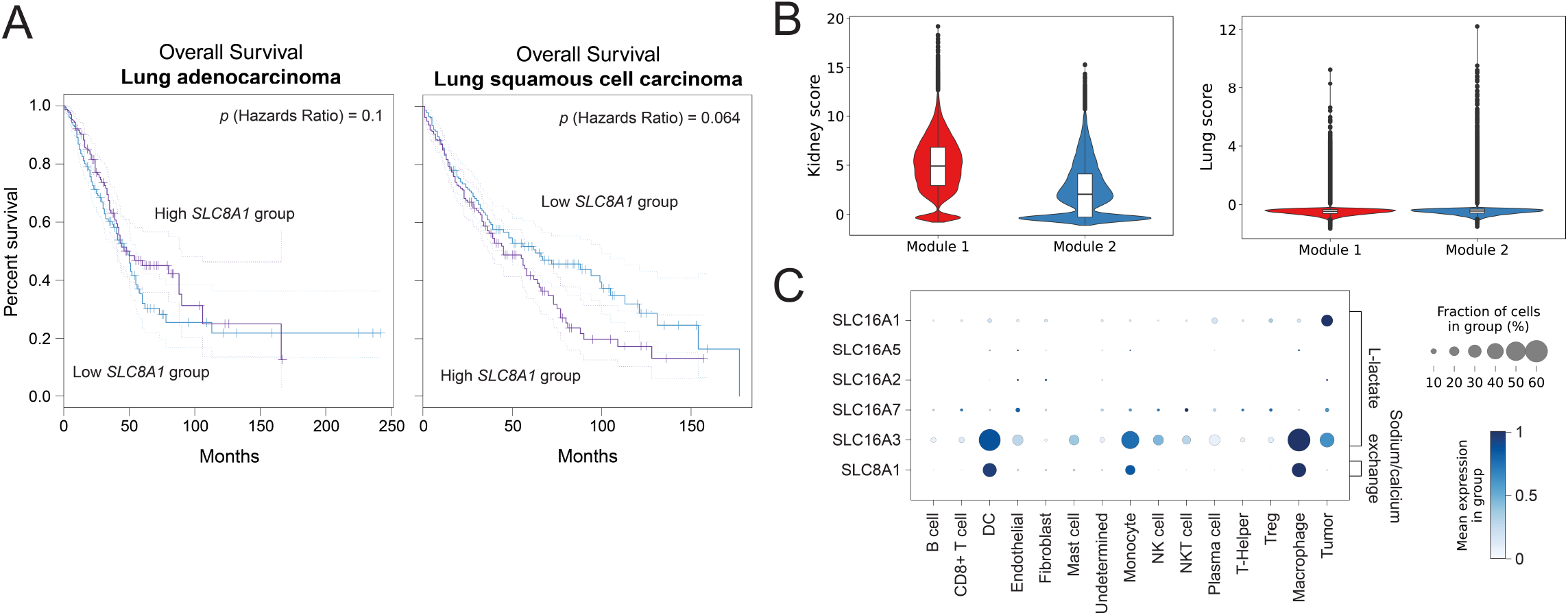
(A) Kaplan-Meier curve of the survival analysis between the *SLC8A1*-high and low groups for lung adenocarcinoma (left) and lung squamous cell carcinoma (right) patients. (B) Violin plots of the kidney and lung signature scores across spots belonging to Modules 1 and 2. (C) Dot plot showing the gene expression of L-lactate and sodium/calcium exchange transporters across cell types from the single-cell dataset used as a reference for deconvolution.

**Figure S7:**
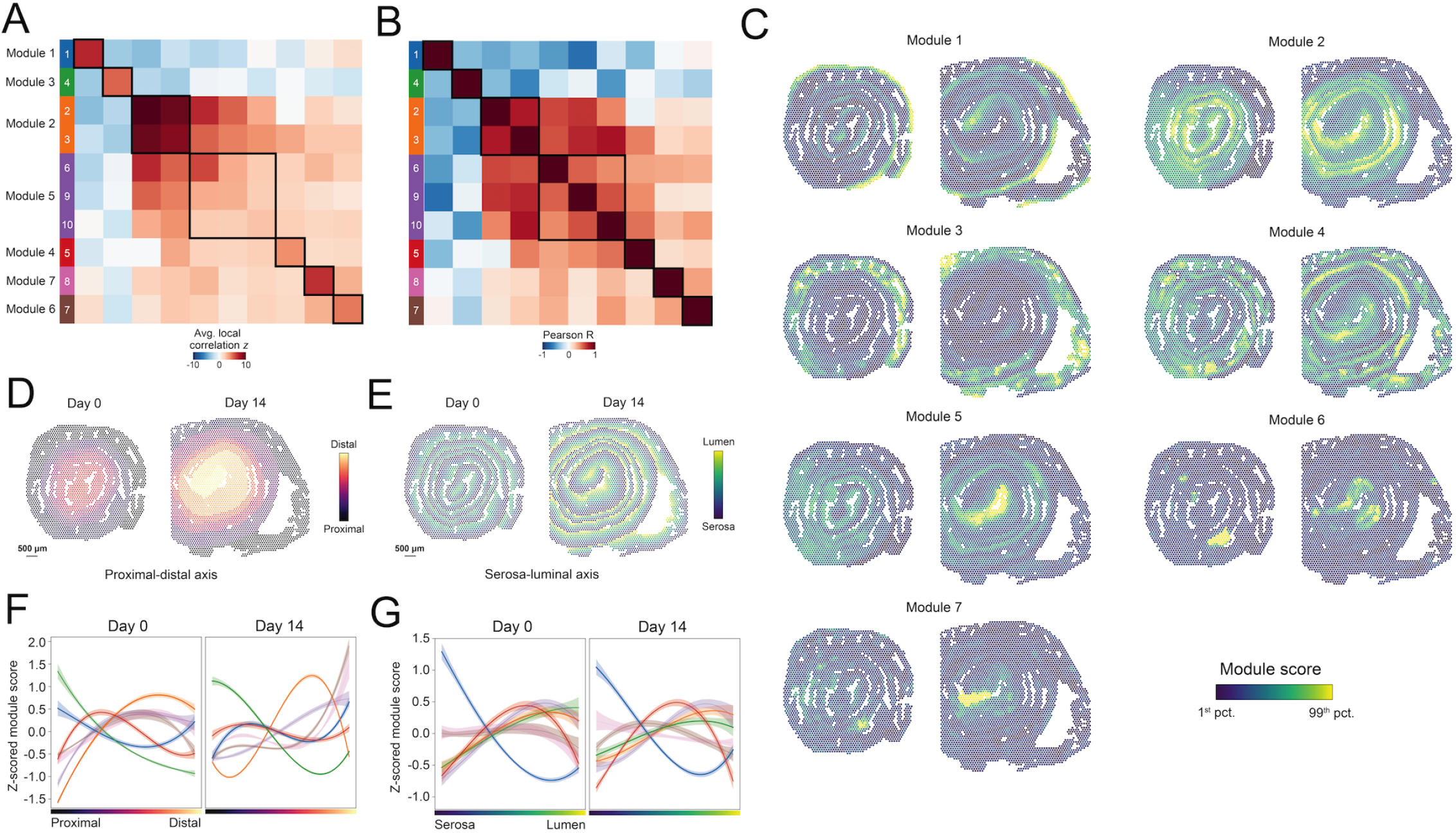
Metabolic zonation on the mouse colon dataset. (A) Cluster map of the average pairwise local correlation (**Test statistic 2**) Z-score between all pairs of genes assigned to the corresponding metabolic gene module pair (row and column) before grouping. Numbered squares are colored, and rows are labeled according to the final module annotation after grouping. (B) Cluster map of the Pearson correlation between the module scores of all pairs of metabolic gene modules (row and column) before grouping. Numbered squares are colored according to the final module annotation after grouping. (C) Spatial plots of the metabolic gene module scores after grouping. (D) Spatial plot colored by the proximal-distal axis. Scale bars, 500 *µ*m. (E) Spatial plot colored by the serosa-luminal axis. Scale bars, 500 *µ*m. (F) and (G) Line plots showing the Z-scored module scores along the proximal-distal (F) and serosa-luminal (G) axes separately for each condition. Scores were smoothed using a polynomial regression model, and mean and 95% confidence intervals (shaded area) are shown. The size of the regression line is proportional to the abundance of a given module in a given condition.

**Figure S8:**
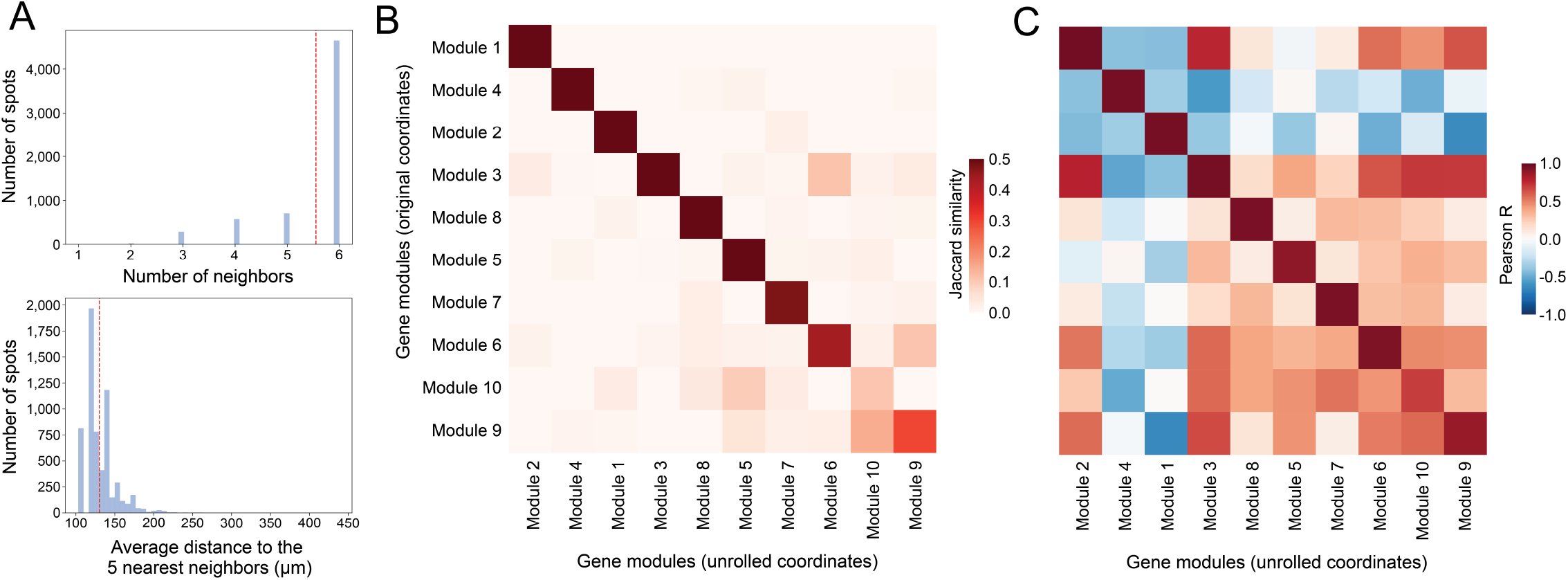
Comparison between the modules computed on the original and the unrolled tissue coordinates. (A) Top: Distribution of the number of neighbors in a neighborhood radius smaller than 150 *µ*m in the original coordinate space. Bottom: Distribution of the average distance to the 5 nearest neighbors in the unrolled space. The vertical dashed red line represents the mean value. (B) Jaccard similarity between the gene modules computed using the original (neighborhood radius of 150 *µ*m) and the unrolled (5 nearest neighbors) coordinates. (C) Pearson correlation between the module scores computed using the original (neighborhood radius of 150 *µ*m) and the unrolled (5 nearest neighbors) coordinates.

**Figure S9:**
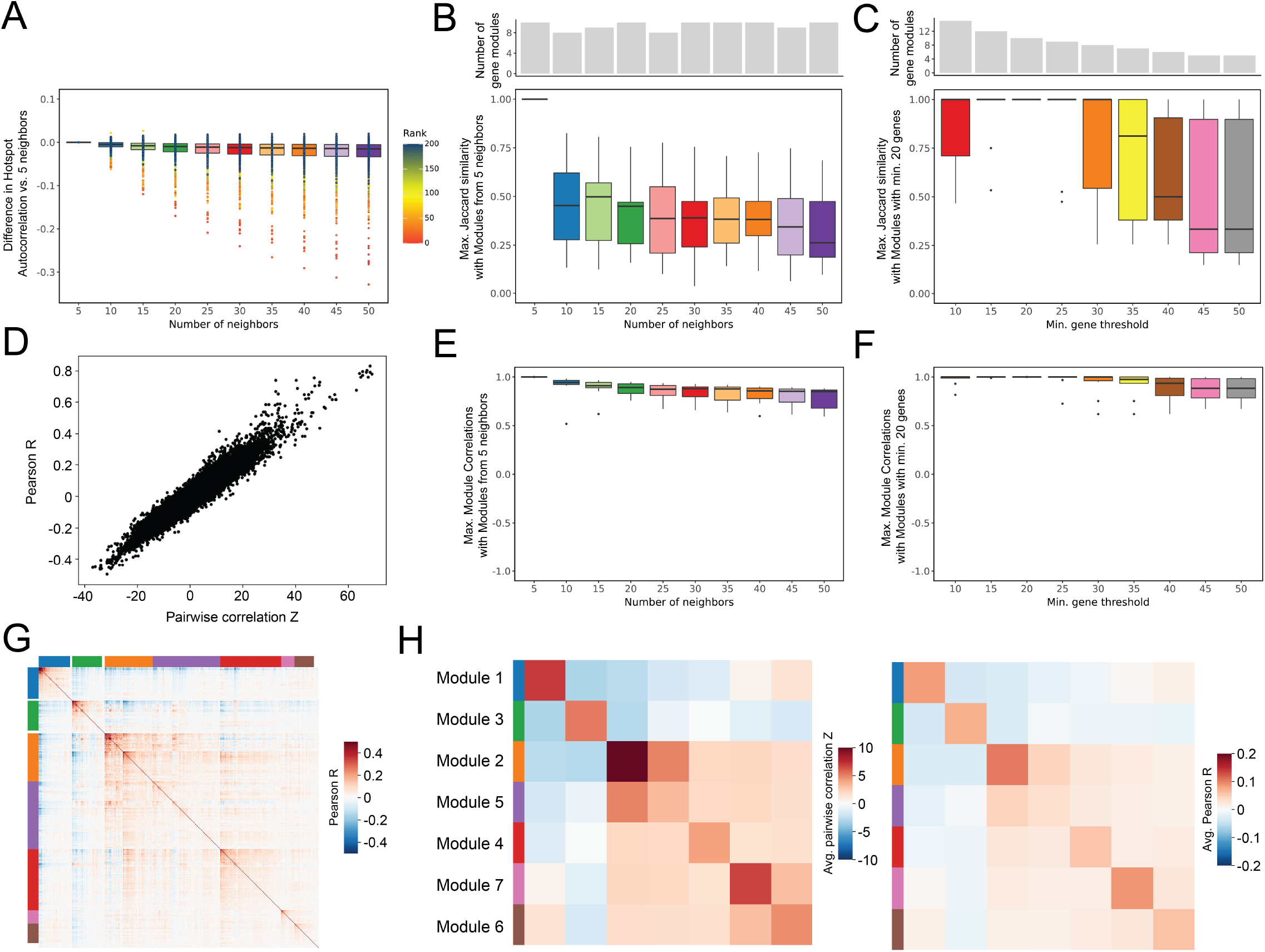
Gene autocorrelation and detected modules in the mouse colon dataset are robust to varying parameter values. (A) Difference in gene autocorrelation (**Test statistic 1**) score between 5 neighbors and a varying number of neighbors. (B) Top: Number of gene modules across a varying number of neighbors. Bottom: Jaccard similarity between the inferred modules in this dataset (5 neighbors) and the most similar modules generated using a different number of neighbors. (C) Top: Number of gene modules across varying minimum gene threshold values. Bottom: Jaccard similarity between the inferred modules in this dataset (minimum gene threshold of 20 genes) and the most similar modules generated using a different minimum gene threshold value. (D) Comparison of Pearson correlation R and pairwise correlation (**Test statistic 2**) Z values for all significantly autocorrelated (theoretical FDR < 0.01; **Test statistic 1**) genes. (E) Pearson correlation between the inferred modules in this dataset (5 neighbors) and the most similar modules generated using a different number of neighbors. (F) Pearson correlation between the inferred modules in this dataset (minimum gene threshold of 20 genes) and the most similar modules generated using a different minimum gene threshold value. (G) Gene-gene Pearson correlation for the 759 genes selected using **Test statistic 1**. (H) Average pairwise correlation Z (left; **Test statistic 2**) and Pearson R (right) values.

**Figure S10:**
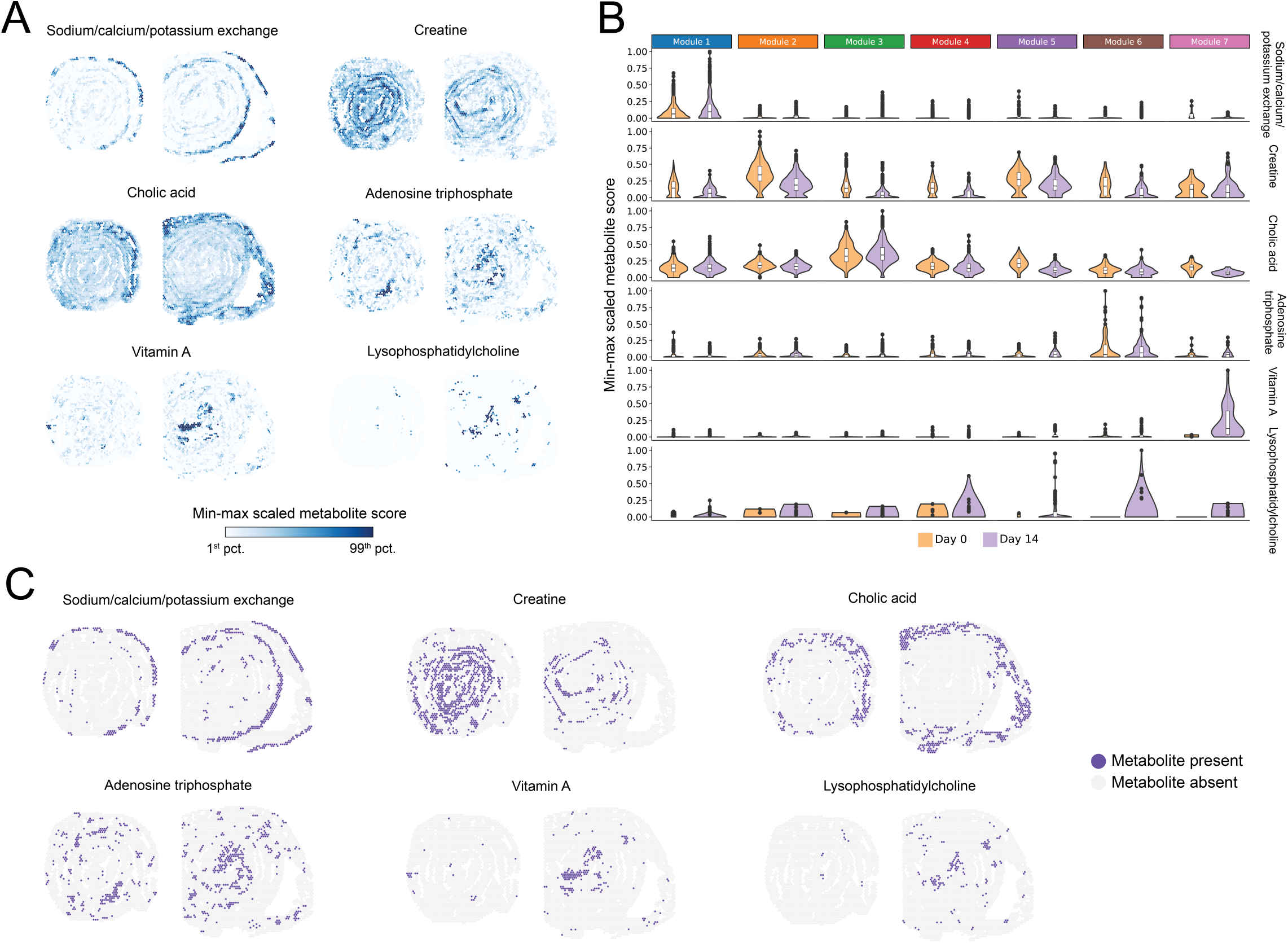
Metabolite abundance across conditions in the mouse colon dataset. (A) Spatial plots of the min-max scaled metabolite scores (**Test statistic 8**) for region- and condition-specific metabolites. (B) Violin plots of the distribution of min-max scaled metabolite scores across modules and conditions. (C) Spatial plots of the binarized metabolite scores used for metabolite module grouping (**Test statistic 4**). Those scores with a Z-scored value higher than 1 standard deviation above the mean were labeled as present.

## Supplementary Tables

**Supplementary Table 1:**
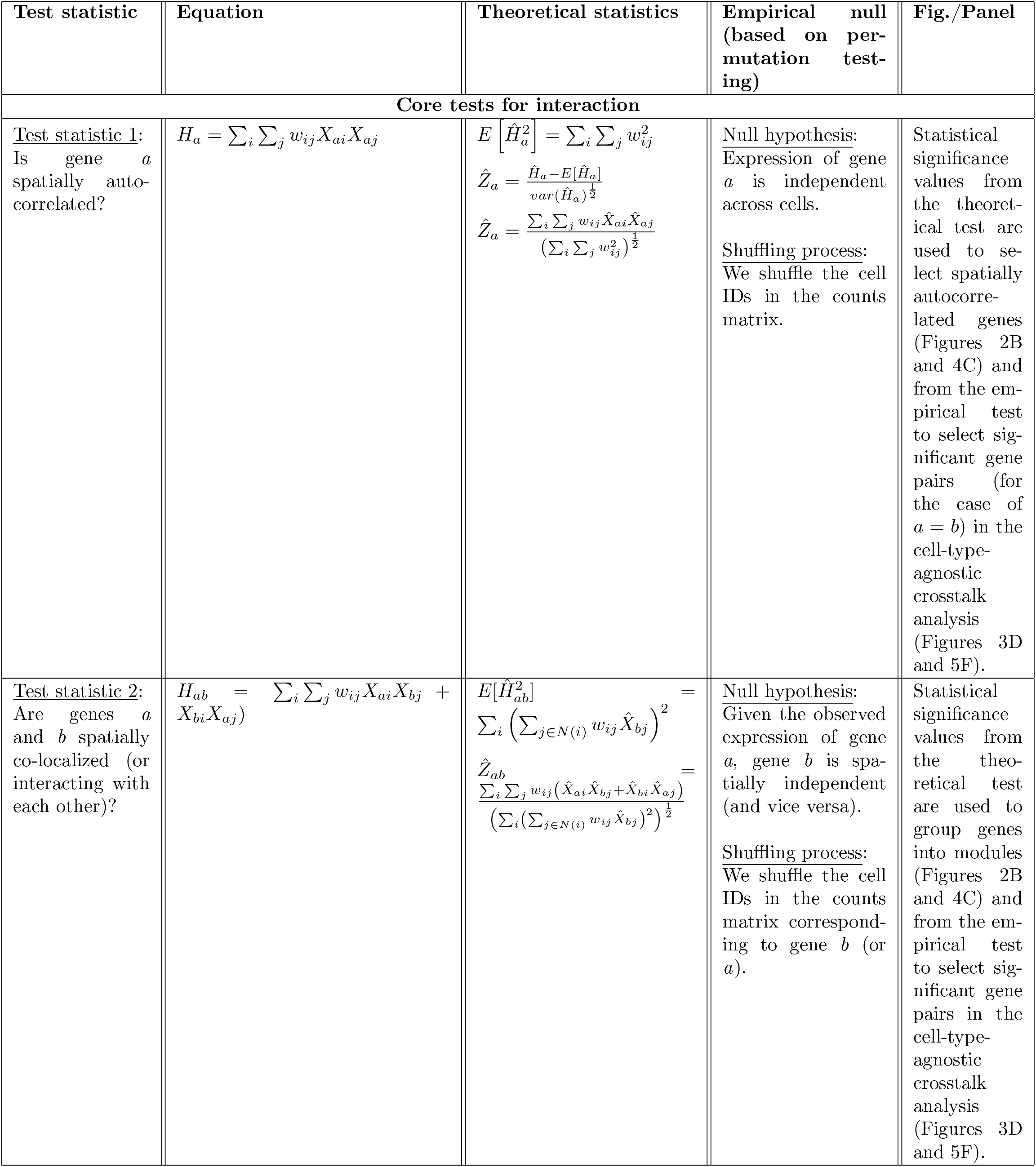
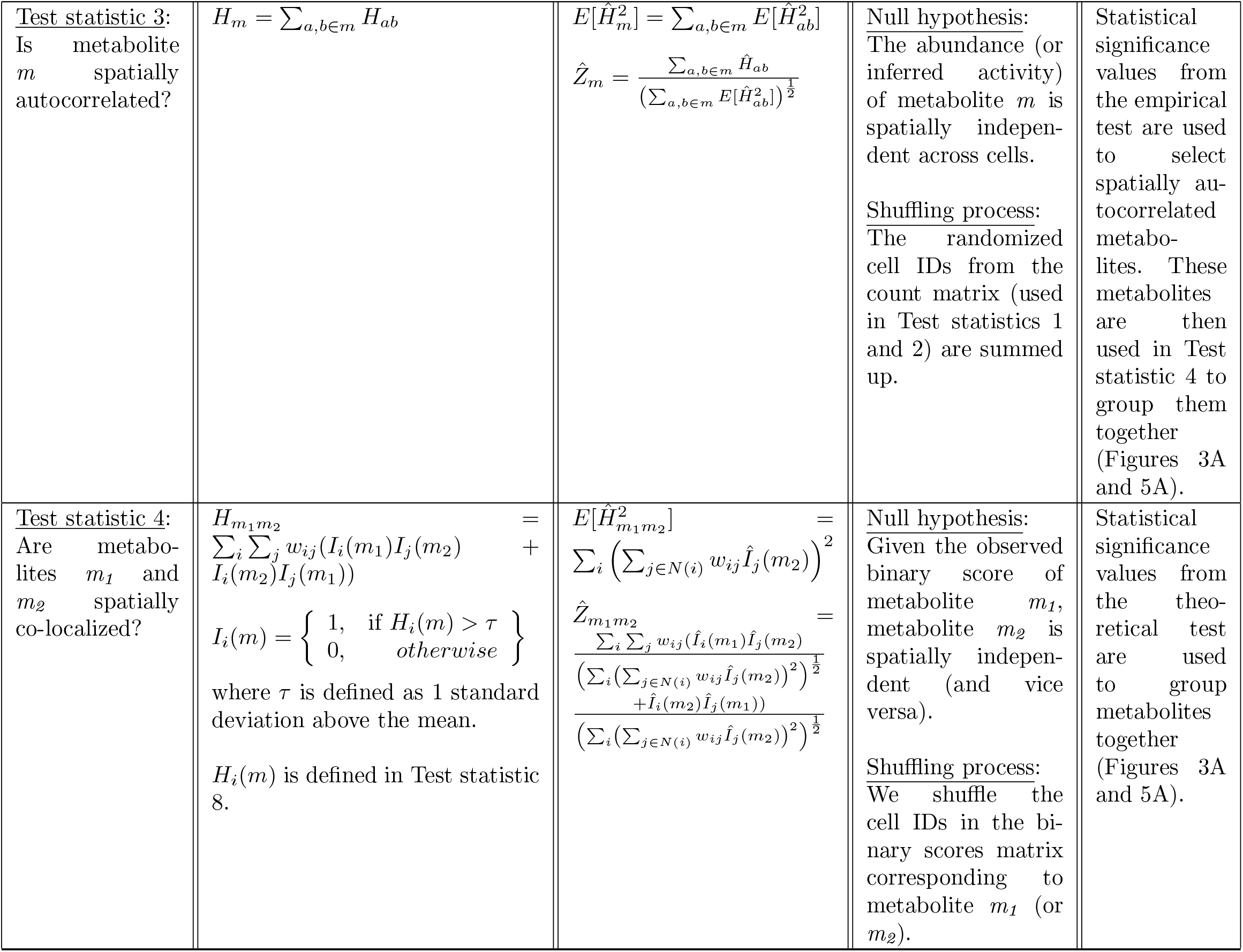
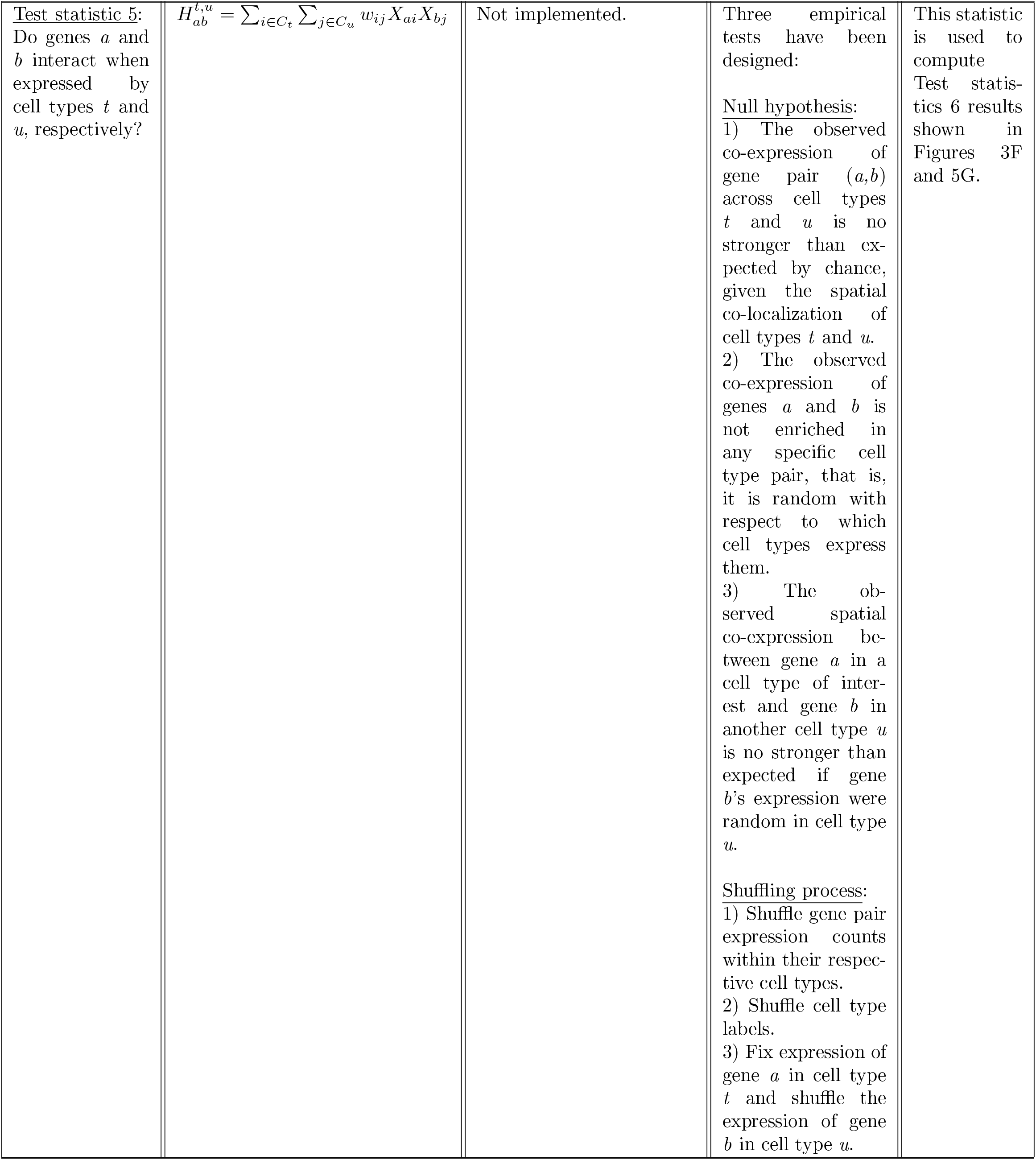
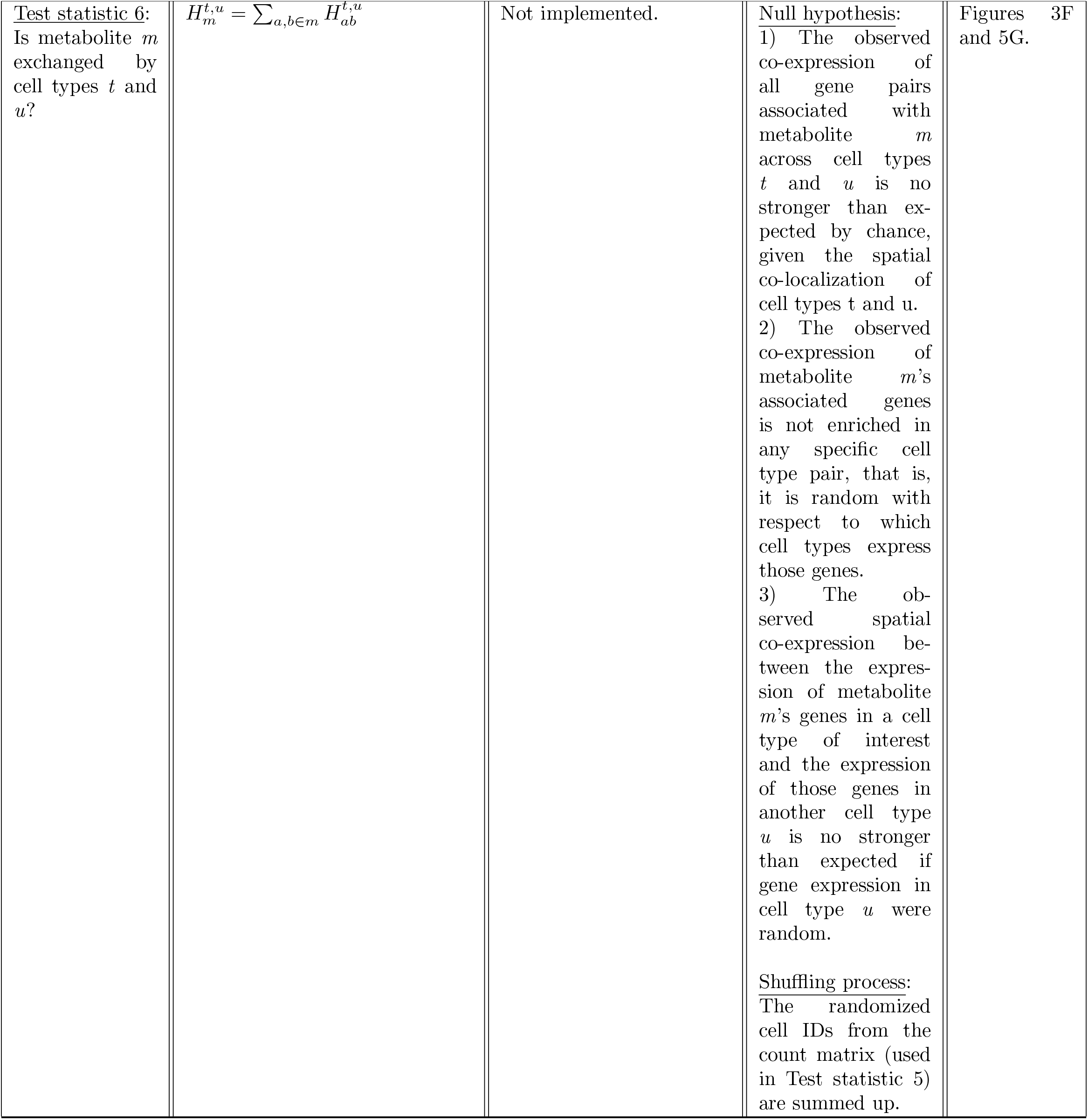
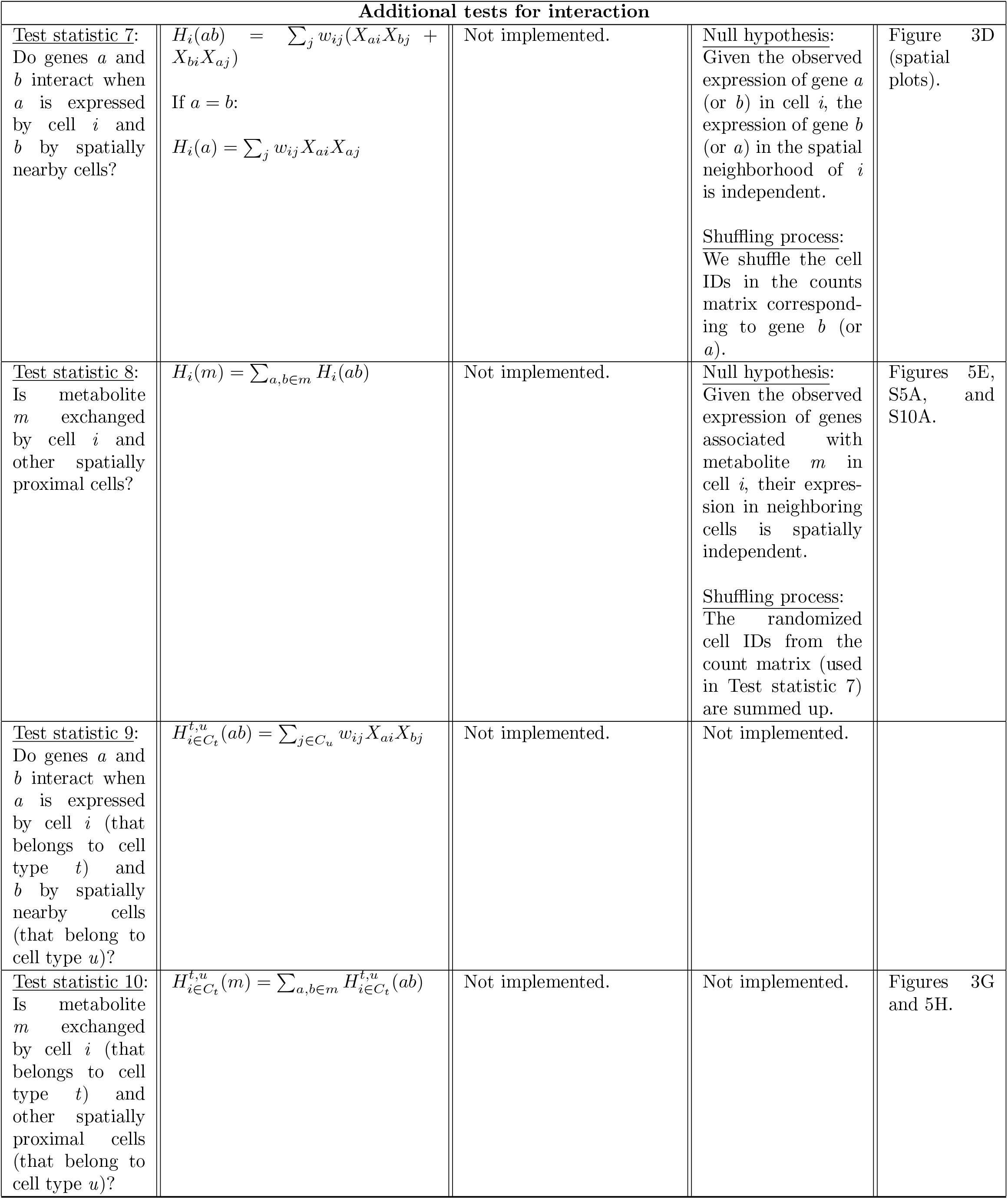
Summary of Harreman test statistics.

**Supplementary Table 2:**
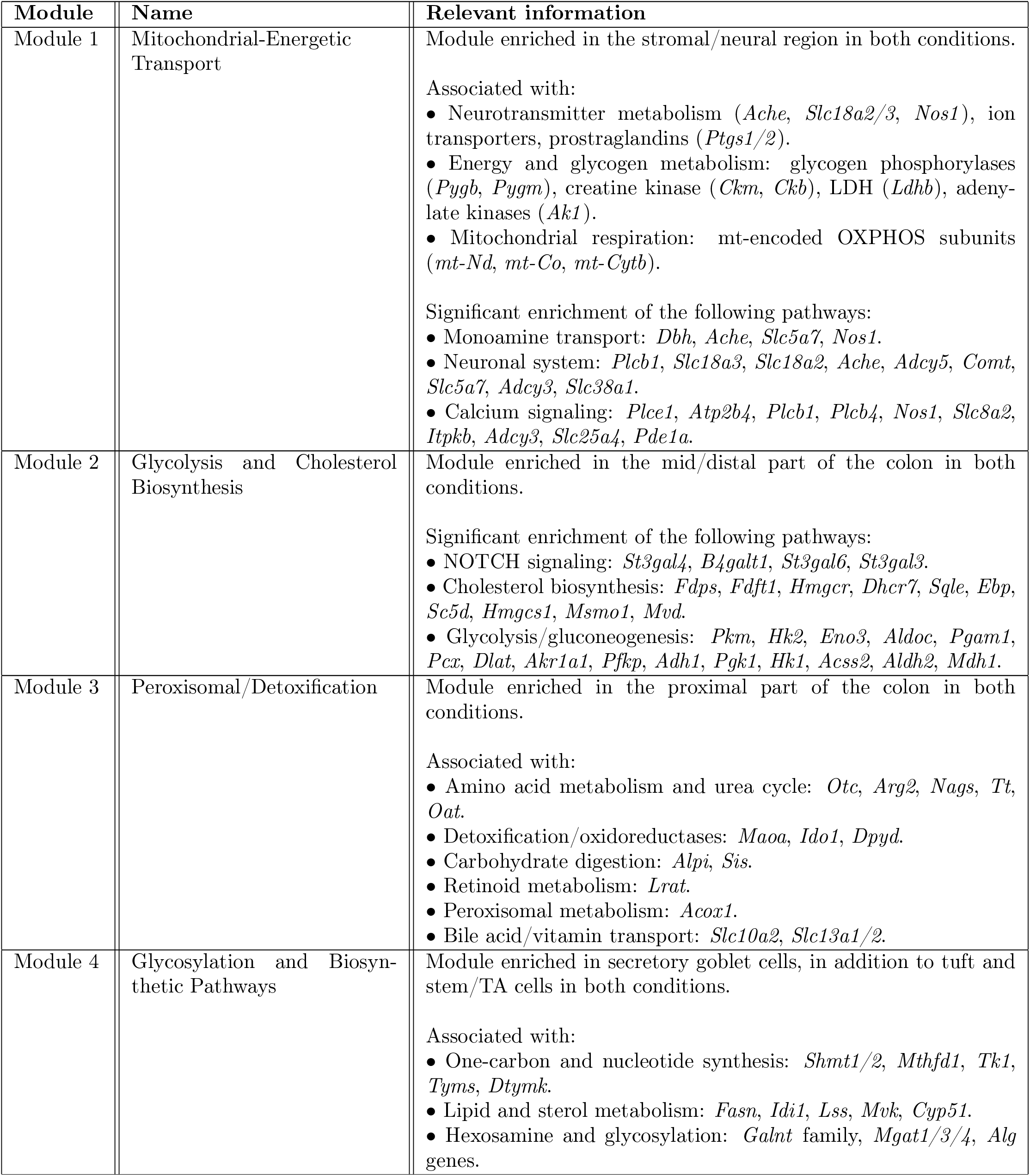
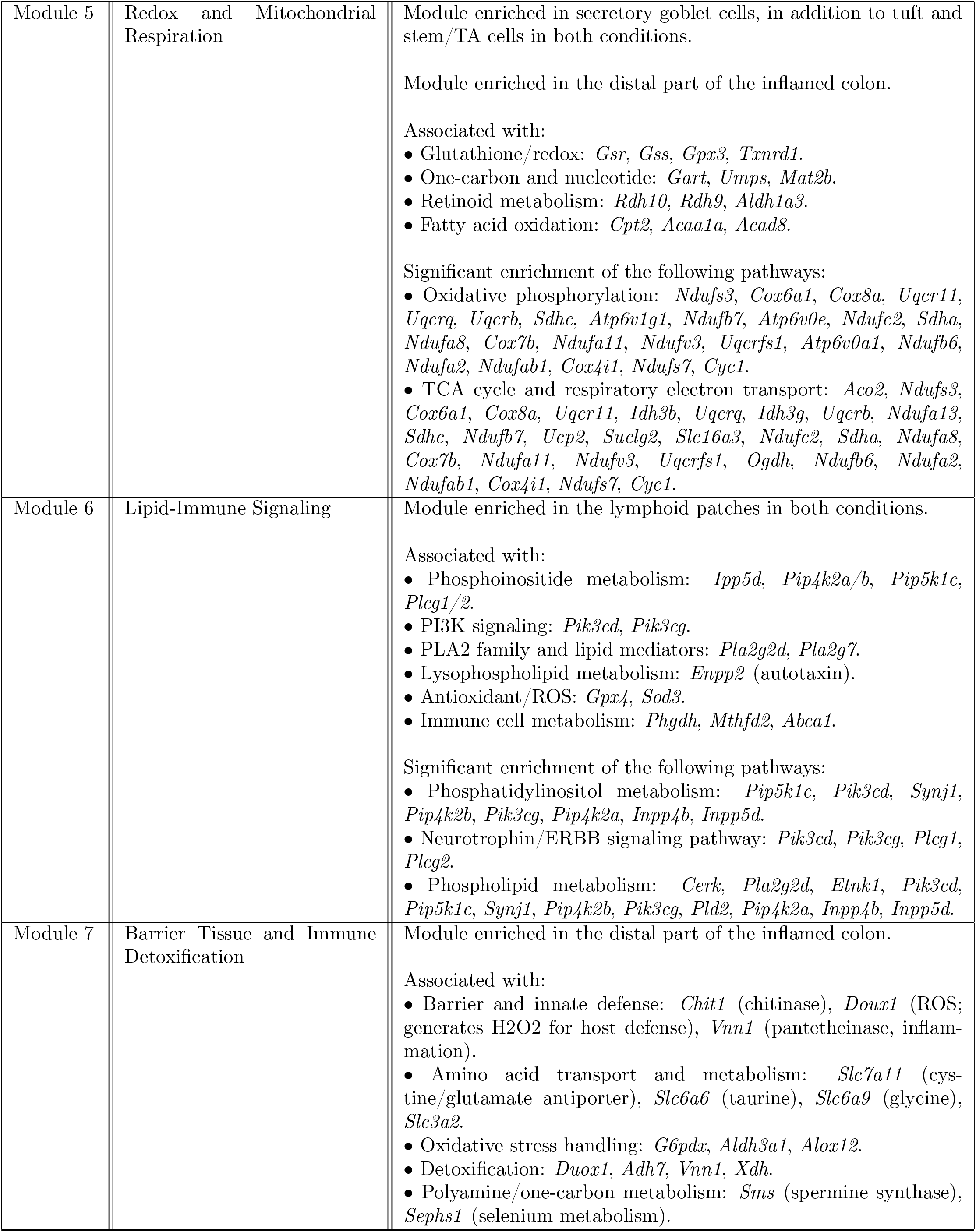
Description of the metabolic modules from the mouse colon dataset.

## Notes

### Competing Interest Statement

The authors have declared no competing interest.

